# miRNA408 and its encoded peptide, miPEP408, regulate arsenic stress response in *Arabidopsis*

**DOI:** 10.1101/2022.04.27.489731

**Authors:** Ravi Shankar Kumar, Hiteshwari Sinha, Tapasya Datta, Mehar Hasan Asif, Prabodh Kumar Trivedi

## Abstract

MicroRNAs (miRNAs) are small non-coding RNAs that play a central role in regulating various developmental and biological processes. The expression of miRNAs is differentially modulated in response to various stresses. Based on the recent findings, it has been shown that some of the pri-miRNAs encode small regulatory peptides, microRNA-encoded peptides (miPEP). miPEPs are reported to regulate the growth and development of plants by modulating corresponding miRNA expression; however, the role of these peptides in different stresses has not been explored yet. Here, we reported that pri-miR408 encodes a small peptide, miPEP408, that regulates the expression of miR408, its targets, and associated phenotype in *Arabidopsis*. Plants overexpressing miR408 showed severe sensitivity under low sulphur (LS), Arsenite As(III) and LS+As(III) stress, while miR408 mutant developed through the CRISPR/Cas9 approach showed tolerance. Transgenic lines showed phenotypic alteration and modulation in the expression of genes involved in the sulphur reduction pathway and affect sulphate and glutathione accumulation. Similar to miR408 overexpressing lines, the exogenous application of synthetic miPEP408 or miPEP408 overexpression led to sensitivity in plants under LS, As(III) and combined LS+As(III) stress compared to control. This study suggests the involvement of miR408 and miPEP408 in heavy metal and nutrient deficiency responses.

**One-sentence summary:** miR408 and peptide encoded by miR408, miPEP408, regulate arsenic stress and low sulphur responses in Arabidopsis.

## INTRODUCTION

MicroRNAs (miRNAs) are small endogenous, non-coding RNAs that act on their targets and down-regulate them through translational repression or mRNA cleavage (Tiwari et al., 2014; Meyers and Axtell, 2019; Li and Yu, 2021). Therefore, these miRNAs control many developmental and physiological processes and are key regulators of plant growth and development (Millar et al., 2020; Shriram et al., 2016; Singh et al., 2016). Several studies have shown that miRNA has an important regulatory role in biotic and abiotic stresses, such as nutrient deficiency conditions, which inhibit plant growth and development (Pagano et al., 2021; Basso et al., 2019; Fujii et al., 2005; Lin et al., 2013; Misra et al., 2010; Sharma et al., 2014). Research on miRNA-encoded peptides (miPEPs) began recently since a study revealed that a few pri-miRNAs have one or more open reading frames (ORFs), which regulate the associated miRNAs by enhancing their transcription (Lauressergues et al., 2015; Couzigou et al., 2015; Sharma et al., 2020; Lauressergues et al., 2022; Badola et al., 2022). Although the role of these miPEPs has been explored in various plants (Chen et al., 2020; Couzigou et al., 2016; Badola et al., 2022; Lauressergues et al., 2022) however, their mode of action under different stresses remains unexplored.

Heavy metal stress and nutrient limitation are major concerns across the globe that drastically impairs plant growth and development. Among the several heavy metals known so far, arsenic (As) is a naturally occurring toxic metalloid found in water, soil and rocks. Both natural and anthropogenic activities like mining, industrialization and pesticide raise As contamination in the food chain (Kumar et al., 2015; Abbas et al., 2018). The effects of As include skin cancer in humans whereas, in plants such as Rice, it accumulates in the grains, eventually making the crop unsuitable for consumption. Arsenite [As(III)] and arsenate [As(V)] are two predominant inorganic forms of As that are readily interconvertible (Kumar et al., 2019). As detoxification is most commonly known to be alleviated by Sulphur (S)-mediated detoxification in plant cells (Kumar et al., 2018). Sulphur being an important macronutrient, not only leads to the synthesis of the two amino acids cysteine and methionine, but also produces glutathione (GSH) and phytochelatins (PCs) essential in combating As stress (Dixit et al., 2015; 2016). Sulphate transportation and assimilation are regulated by several modes, including transcriptional and post-transcriptional regulation in plants. Transcriptional regulation is mediated by factors such as sulphate limitation, light, sugars, heavy metals, etc., whereas sulphate assimilation is post-transcriptionally regulated by miR395 (Yuan et al., 2016; Li et al., 2020). Interestingly, a study revealed the differential modulation in the expression of various miRNAs, among which miR408 was identified to be suppressed in response to sulphate deficiency and carbon and nitrogen limitation (Liang et al., 2015).

miR408 is a highly conserved miRNA and its expression is controlled by the availability of copper (Shahbaz et al., 2019; Kozomara and Griffiths-Jones, 2011; Axtell and Bowman, 2008). The targets of miR408 include genes encoding copper-binding proteins, which belong to the phytocyanin family and function as electron transfer shuttles between proteins (Zhang et al., 2017), and laccase, another copper-containing protein involved in oxidative polymerization of lignin (Tobimatsu et al., 2019; Berthet et al., 2012). The expression of miR408 is significantly affected by a variety of developmental and environmental conditions; however, limited information is available related to its biological function. miR408 has been found as an important component of the HY5–SPL7 gene network that mediates the coordinated response to light and copper, further illustrating its central role in the response of plants to their environment (Zhang et al., 2014). Studies also suggest that PIF1-miR408-Plantacyanin Repression Cascade regulates light-dependent seed germination in Arabidopsis (Jiang et al., 2021). Previous studies illustrated the important role of miR408 in governing tolerance towards abiotic factors such as salinity, cold and oxidative stress, and vice-versa in case of drought and osmotic stresses (Hajyzadeh et al., 2015; Bai et al., 2018; Rajwanshi et al., 2014; Ma et al., 2015). Although the actual mechanism by which miRNA408 regulates different abiotic stresses is not well understood.

This study emphasizes the role of miR408 and the peptide encoded by miR408 (miPEP408) in response to different stresses such as heavy metal and nutrient deficiency. For a better understanding of the role of miR408, overexpressing and CRISPR-edited plants of miR408 were developed. The overexpression plant of miR408 showed a similar phenotype as peptide supplemented plants with increased root length and fresh weight. Though the accumulation of miR408 was significantly decreased and its targets were upregulated in miR408 mutant plants, these did not show a significant phenotypic difference compared to the WT. Interestingly, under sulphate deficit and As stress conditions, miR408-OX lines showed a more sensitive phenotype than the WT. Interestingly, CRISPR-edited miR408 plants exhibited increased root length and fresh weight under stress conditions compared to the WT. Application of synthetic miPEP408 exogenously led to upregulation of miR408 expression that negatively regulates the target genes, Plantacyanin (ARPN) and Laccase3 (LAC3), affecting plant growth in the control condition. Further, miPEP408 supplemented seedlings under different stress conditions showed a highly sensitive phenotype, similar to the miPEP408-OX lines. Therefore, this study gives a wide picture of the response of miR408 towards sulphate limitation and As stress.

## RESULTS

### Differentially regulated miRNAs under low sulphur and arsenic stress

Identification of miRNAs in response to As and sulphur deficiency could be very helpful to decipher the regulatory pathway involving specific miRNAs in these stresses. Thus, to identify miRNAs with modulated expression in response to As stress and sulphur limitation (LS), small RNA sequencing of seedlings grown on C, LS, As(III) and LS+As(III) was performed (NCBI Bioproject: PRJNA830335). The analysis of the small RNA sequencing was done on the basis of the total reads obtained for each sample, raw reads sequence length, total data size obtained and raw reads of samples after adapter removal (Supplemental Table S1 and S2). The total numbers of known miRNAs found in each of the conditions are represented in Supplemental Table S3. Further analysis revealed that 55 miRNAs are differentially regulated under As(III) and LS and LS+As(III) conditions compared to control (Figure 1A; Supplemental Table S4). Out of total differentially expressed miRNAs, 18 miRNAs are commonly differentially expressed under As(III) and LS and LS+As(III) conditions (Figure 1A and 1B; Supplemental Table S5). We identified several miRNAs (miRNA398, miR399, miR408 and miR857), which are down-regulated under As and LS conditions (Figure 1B), however, some of the miRNAs were differentially expressed under specific stress conditions. This indicates that the regulatory effect of a miRNA can vary under different stress conditions and physiological requirements of plants are affected by different stress conditions. Validation of expression of down-regulated miRNAs under different stress conditions suggested that miRNA408 was the most responsive towards As and LS conditions (Supplemental Figure S1).

**Figure 1:**
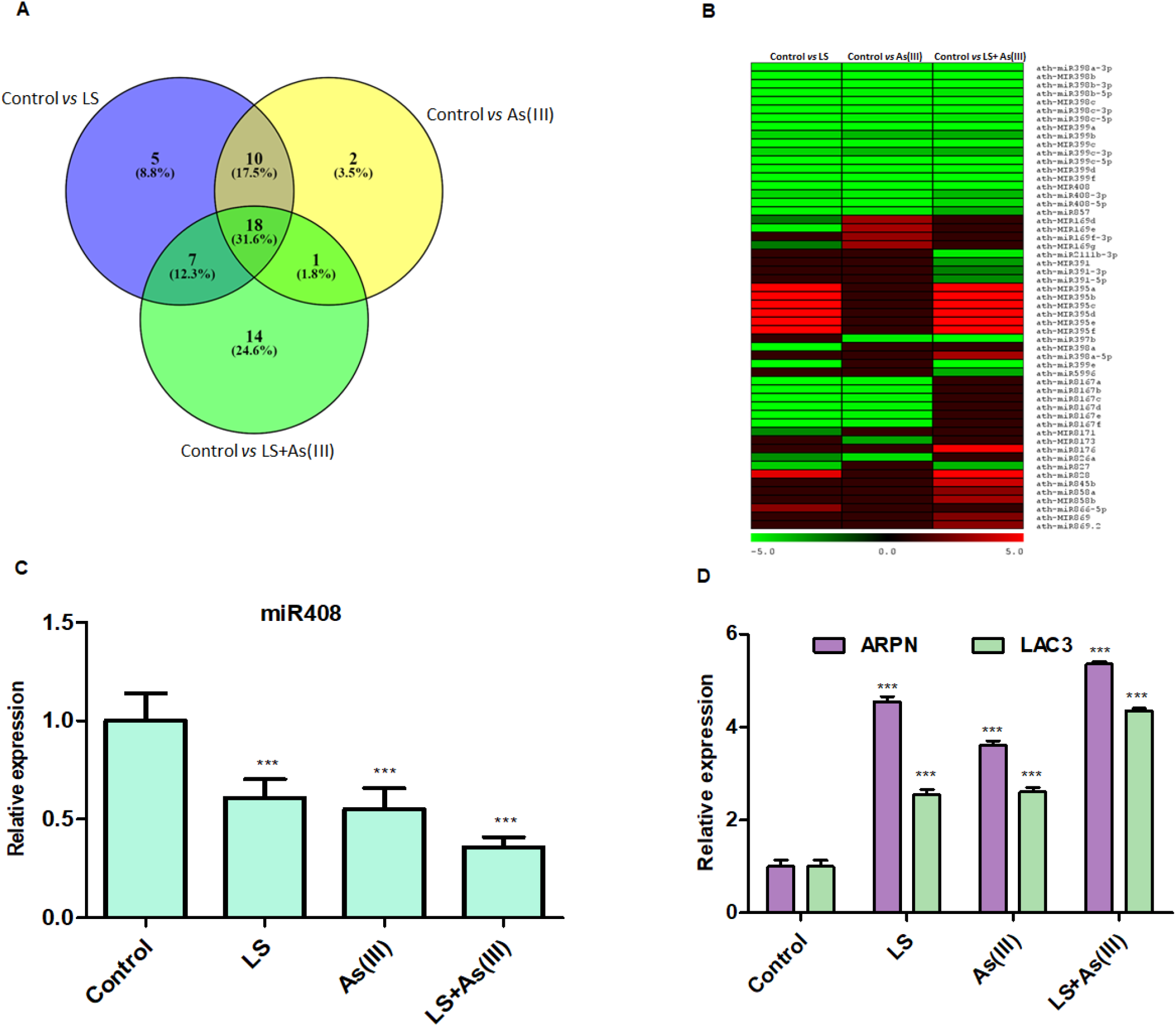
Expression analysis of differentially regulated miRNAs in LS, As(III) and LS+As(III), compared to control. **(A)** Venn diagram of Control Vs LS, Control Vs As(III), Control Vs [LS+As(III)] **(B)** Expression profiles of miRNAs differentially modulated under LS, As(III) and [LS+As(III)]stress in Col-0. The colour scale for fold change values is shown above **(C)** Analysis of pre-miR408 expression in control, LS, As(III) (10 µm) and LS supplemented with As(III) (10 µm). **(D)** Expression analysis of targets ARPN and LAC3 in control, LS, As(III) (10µm) and LS supplemented with As(III) (10 µm). Venn diagram with pvalue <0.05, log2fold change between 1 and −1. Arabidopsis (Col-0) Seeds were grown on C, LS, As(III) and LS+As(III) stress for ten days. Data represent mean ± SD. *** indicate values that differ significantly from control at P < 0.001, according to the student’s unpaired t-test.

The function of miR408 has been studied under plant development and abiotic stresses; however, its role in nutrient deficiency and heavy metal stress needs to be explored. To functionally dissect the role of miR408 towards LS and As stress, we first examined the transcript abundance of miR408 in response to LS and As(III) and combined stress [LS+As(III)]. The expression analysis revealed that miR408 accumulation is reduced under LS and AsIII stress, while the transcript level was lowest under [LS+As(III)] (Figure 1C). Previously, ARPN and LAC3 genes were identified as cleavable targets of miR408 (Abdel-Ghany and Pilon, 2008). Therefore, we analysed the expression of *ARPN* and *LAC3* in the above stress condition. Based on expression analysis, it was observed that LS, As(III) and [LS+As(III)] stress conditions influence the *ARPN* and *LAC3* expression in a pattern contrary to that of miR408 (Figure 1D). These results indicate that miR408 is regulated by LS and As(III) stress and modulates target expression under these conditions.

### *MIR408* is transcriptionally regulated by LS and As

To investigate the promoter responsiveness towards LS, As and [LS+As(III)] stress, promoter-reporter lines (wild-type seedling expressing *promiR408::GUS*) and positive control [Empty vector (EV) with GUS gene under CaMV35S] were grown under different stress conditions. The GUS histochemical analysis suggests that compared to the EV, *promiR408::GUS* lines showed much reduced GUS activity under LS, As(III) and [LS+As(III)] stress conditions (Figure 2A), consistent with the miR408 transcript analysis (Figure 2B). To correlate the promoter activity with miR408 expression, the miR408 expression was analysed in all promoter-reporter lines. The analysis revealed significantly decreased transcript levels of miRNA408 in LS, As(III) and [LS+As(III)] (Figure 2C). The target gene expression analysis for miR408 (APRN) showed a higher accumulation under stress conditions than in control (Figure 2D). These results suggest that the promoter activity and transcript level of miRNA408 are significantly affected by LS and AS stress conditions.

**Figure 2.**
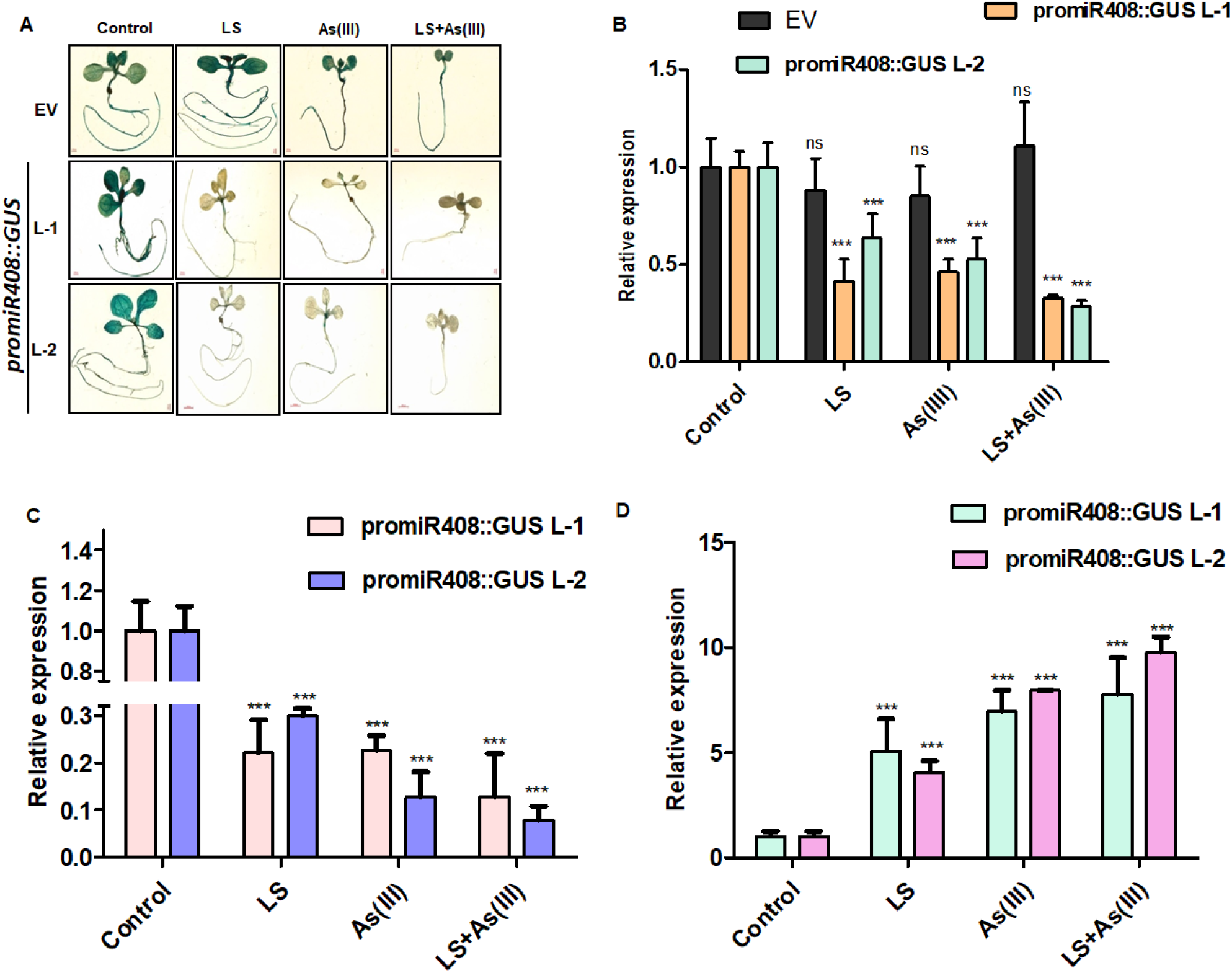
miR408 Participates in the Coordinated Low Sulphur (LS) and Arsenite As(III)) response. (**A)** Histochemical GUS staining in ten-day-old Arabidopsis seedlings transformed with EV (CaMV 35S::GUS) and Pro:miR408::GUS using L1 and L2 lines on control LS, As(III) and combined [LS+As(III)] conditions. Scale bars, 1,000 μm. **(B)** Expression analysis of GUS in 10d old seedlings transformed with EV and Pro:miR408::GUS using L1 and L2 lines on control LS, As(III) and combined [LS+As(III)] conditions using quantitative-RTPCR. (**C)** Analysis of pre-miR408 expression in control, LS, As(III) (10 µm) and LS supplemented with As(III) (10 µm) using Pro:miR408::GUS lines (**D)** Expression analysis of targets ARPN in control, LS, As(III) (10µm) and LS supplemented with As(III) (10 µm) using Pro:miR408::GUS lines. The experiment was repeated with three biological replicates with similar results. The student’s t-test was used for statistical analysis and error bars were represented as three technical replicates. Tubulin was used as an internal control to quantify the relative expression of the gene.

### Responses of miR408 overexpression and CRISPR/Cas9-based knockout mutants towards As and LS

To study the impact of LS and As stresses on miR408, miR408 overexpressing lines (mi408OX) and mutated plants (*miR408^CR^*) were developed (Supplemental Figure S2A **and** S2B). Three homozygous miR408OX lines were selected on the basis of enhanced levels of miR408 expression and decreased expression of target genes (*ARPN, LAC3*) compared to WT (Figure 3A and 3C). To develop miR408 mutant plants, we used CRISPR/Cas9 approach, in which gRNA was designed from the pre-miRNA region targeting mature miRNA. However, screening of mutants by sequence analysis revealed deletion and insertions in the mature miRNA sequences obtained (Supplemental Figure S3A **and** S3B). Three homozygous and Cas9-free edited plants of the T4 generation were used for further study. These mutated plants showed significantly reduced miR408 expression and higher accumulation of target genes (*ARPN*, *LAC3*) (Figure 3B and 3D).

**Figure 3.**
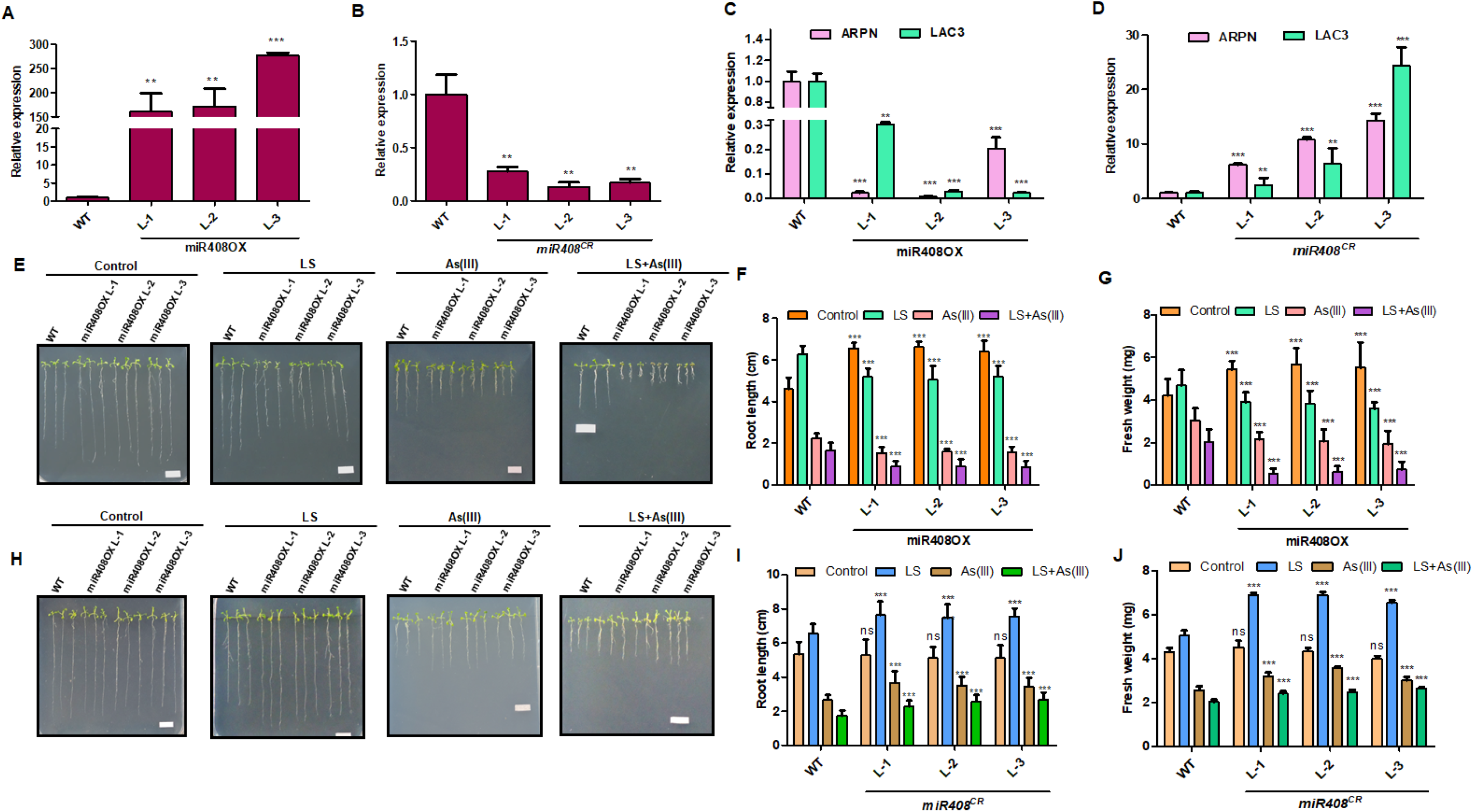
Phenotypic variation of miR408-OX and CRISPR/Cas9 derived knockout mutants towards limiting sulphur and arsenic stress conditions. **(A)** Expression analysis of pre-miR408 in 10 days old WT and miR408-OX plants. Three independent experiments were performed, with similar results. (**B**) Expression analysis of pre-miR408 in 10 days old WT and *miR408^CR^* plants. Three independent experiments were performed, with similar results. (**C)** Relative expression of *ARPN* and *LAC3* in 10 days old WT and miR408 overexpressing plants. (**D)** Relative expression *ARPN* and *LAC3* in WT and in 10 days old seedlings. (**E)** Representative image of ten-day-old WT and miR408-OX seedlings grown on control, LS, As(III) and [LS+As(III)] conditions. Scale bar, 1 cm. **(F**) Primary Root lengths of ten-day-old WT and miR408-OX seedlings grown on control, LS, As(III) and [LS+As(III)] conditions. N=30 independent seedlings. (**G)** Fresh weight of ten-day-old WT and miR408-OX seedlings grown on control, LS, As(III) and [LS+As(III)] conditions. N=30 independent seedlings. (**H**) Representative image of ten-day-old WT and *miR408^CR^* seedlings grown on control, LS, As(III) and [LS+As(III)] conditions. Scale bar, 1 cm. (**I)** Primary Root lengths of ten-day-old WT and *miR408^CR^* seedlings grown on control, LS, As(III) and [LS+As(III)] conditions. N=30 independent seedlings. (**J)** Fresh weight of ten-day-old WT and *miR408^CR^* seedlings grown on control, LS, As(III) and [LS+As(III)] conditions. N=30 independent seedlings. Calculations of data were performed from three biological replicated independently per treatment with similar results. The asterisk denotes significant difference in values with *** as *P < 0.1; **P < 0.01; ***P < 0.001, according to two-tailed student’s t-test.

To functionally validate the role of miRNA408 in LS and As stress, miR408OX lines and *miR408^CR^* plants were grown under LS, As(III) and combined [LS+As(III)] stress. Phenotypic analysis of miR408OX lines revealed better growth and root length compared to the WT under control conditions. However, under LS, As(III) and [LS+As(III)] stress, miR408OX lines exhibited retarded growth and a decrease in primary root length compared to WT (Figure 3E and 3F). These stresses affected the plant growth, which finally reduced the fresh weight of miR408OX lines compared to the WT (Figure 3G). On the contrary, compared with WT, the *miR408^CR^* plants were tolerant and exhibited better growth compared to WT under stress conditions (Figure 3H). Moreover, miR408*^CR^* plants have significantly increased primary root length and the fresh weight as compared to the WT plants under stress conditions (Figure 3I and 3J). Together, these results suggest that mutation in miR408 showed tolerance towards LS, As(III) and combine [LS+As(III)] stress. In contrast, the miR408OX lines were sensitive to these stresses compared to the WT. Altogether, a significant change in the phenotypes in miR408OX and *miR408^CR^* suggests that miR408 plays a very important role in nutrient limitation and heavy metal stress.

### miR408 elevates ROS production under LS and As toxicity

Plant cellular machinery increases the production of reactive oxygen species (ROS) under environmental cues such as nutrient deficit, temperature, heavy metal stress etc (Huang et al., 2019; Hasanuzzaman et al., 2020). Higher ROS generation leads to oxidative stress in plants and hampers plant growth. Since miR408OX lines were identified to be sensitive under the different stresses as compared to *miR408^CR^* plants and WT, ROS levels were analysed through NBT and DAB staining. NBT staining revealed that miR408OX lines were accumulating more O^2-^, but the mutated plants showed lesser accumulation compared to the WT. Maximum staining was observed under combined [LS+As(III)] stress, which indicates that mutated lines have decreased oxidative burst as compared to the miR408OX lines thereby leading to sensitivity in them (Figure 4A, Supplemental Figure S4). Furthermore, DAB staining was performed to monitor the production of H_2_O_2_ under LS, As(III) and [LS+As(III)] conditions. Higher accumulation of H_2_O_2_ was observed in miR408OX lines marked as higher brown precipitate. Similar to NBT, maximum DAB staining was found in response to [LS+As(III)] stress, however, *miR408^CR^* plants were observed to accumulate lesser H_2_O_2_ compared to WT (Figure 4B, Supplemental Figure S5).

**Figure 4.**
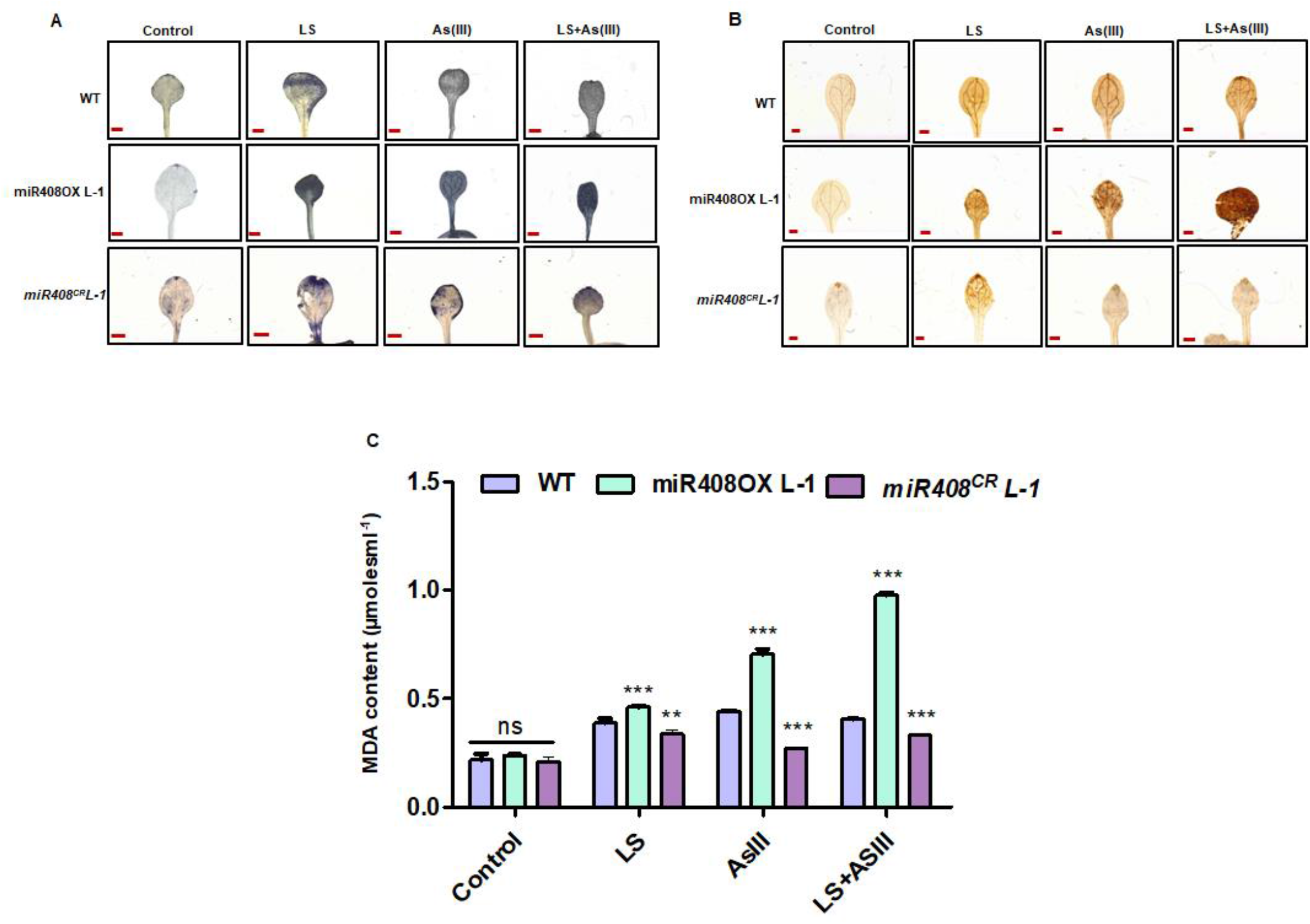
Differential ROS production and lipid peroxidation in miR408-OX and *miR408^CR^* plants in limiting sulphur and As(III) stress. Ten-day-old WT, miR408-OX and *miR408^CR^* were grown for 10 days on medium containing optimum sulphur as control (C), limiting sulphur (LS), As (III) (10 μM), [LS + As (III)] (10μM). (**A)** Staining of WT, miR408-OX (L-1) and *miR408^CR^* (L-1) seedlings with Nitrotetrazolium (NBT). (**B)** Staining of WT, miR408-OX (L-1) and *miR408^CR^* (L-1) seedlings with 3-3’diaminobenzidine (DAB) (**C)** Estimation of MDA in WT, miR408-OX (L-1) and *miR408^CR^* (L-1) seedlings after Control and LS, As(III) and [LS+As(III)] treatment. This data was generated from three biological replicated independently per treatment with similar results. The asterisk denotes significant difference in values with *** as *P < 0.1; **P < 0.01; ***P < 0.001, according to two-tailed student’s t-test. bar=1mm.

Previous studies showed that As(III) induces oxidative stress and generation of free radicals causes lipid peroxidation and tissue damage (Møller et al., 2007; Shri et al., 2009). Malondialdehyde (MDA) content was measured to quantify lipid peroxidation in miR408OX lines and *miR408^CR^* plants after stress treatment. The miR408OX lines showed significantly higher MDA content in LS, As(III) and [LS+As(III)] stress whereas the mutated plants showed lesser accumulation of MDA compared to WT (Figure 4C, Supplemental Figure S6A **and** 6B). These results suggest that the *miR408^CR^* plants have stronger antioxidant and detoxification capability than the miR408OX lines, and was correlated with the phenotypic data, which showed mutated plants are more tolerant compared to miR408OX lines towards LS, As(III) and [LS+As(III)] stress.

### Differential regulation of sulphur metabolism affects detoxification mechanism of miR408OX and *miR408^CR^* plants

Synthesis of various Sulphur (S) containing metabolites, including GSH and amino acids (Cys and Met) is regulated by enzymes involved in sulphate assimilation in plants. This pathway is tightly regulated by the demand and supply of S by the various key factors (Aarabi et al., 2020; Leustek et al., 2000). Previous studies reported that heavy metal stress affects S assimilation in the plant by regulating the expression of key enzymes in the pathway (Khare et al., 2017). So, to correlate the effect of miR408 on S metabolism and As toxicity, we analysed the expression of genes majorly involved in the sulphur reduction in miR408OX as well as *miR408^CR^* plants.

Differential expression of *AtAPS1,2* and *3* were found in miR408OX and *miR408^CR^* plants under LS, As(III) and combined [LS+As(III)] stress comparison to control. Compare to the WT, the miR408OX lines showed a lower accumulation of *AtAPS1, AtAPS2* and *AtAPS3* transcripts, which suggests that there is a significant decrease of flux towards the reduction pathway. Due to this limitation, miR408OX plants might have experienced a limitation of reduced compounds to combat stress (Figure 5A, Supplemental Figure S7A **and** S7B). In plants, heavy metal stress induces the accumulation of GSH via activating ATPS accumulation (Anjum et al., 2015; Asgher et al., 2014), so this pathway needs to be upregulated to combat stress. Contrary to the miR408OX, *miR408^CR^* plants were observed to have a higher accumulation of *AtAPS1, AtAPS2* and *AtAPS3* (Figure 5B; Supplemental Figure 8A and 8B), which suggests the increased flux towards the reduction pathway and higher accumulation of GSH which would be responsible for tolerance of mutated plants under different stresses. The expression of *AtAPR1,2* and *3* in the miR408OX lines was reduced compared to the WT (Figure 5C, Supplemental Figure 7C and 7D), while significantly enhanced expression of APR isoforms was observed in *miR408^CR^* plants in response to LS, As(III) and [LS+As(III)] conditions (Figure 5D, Supplemental 8C and 8D). Enhanced APS and APR activities in *miR408^CR^* plants compared to miR408OX lines might lead to higher accumulation of GSH in *miR408^CR^* plants, therefore, providing tolerance towards LS, As(III) and [LS+As(III)] stress.

**Figure 5.**
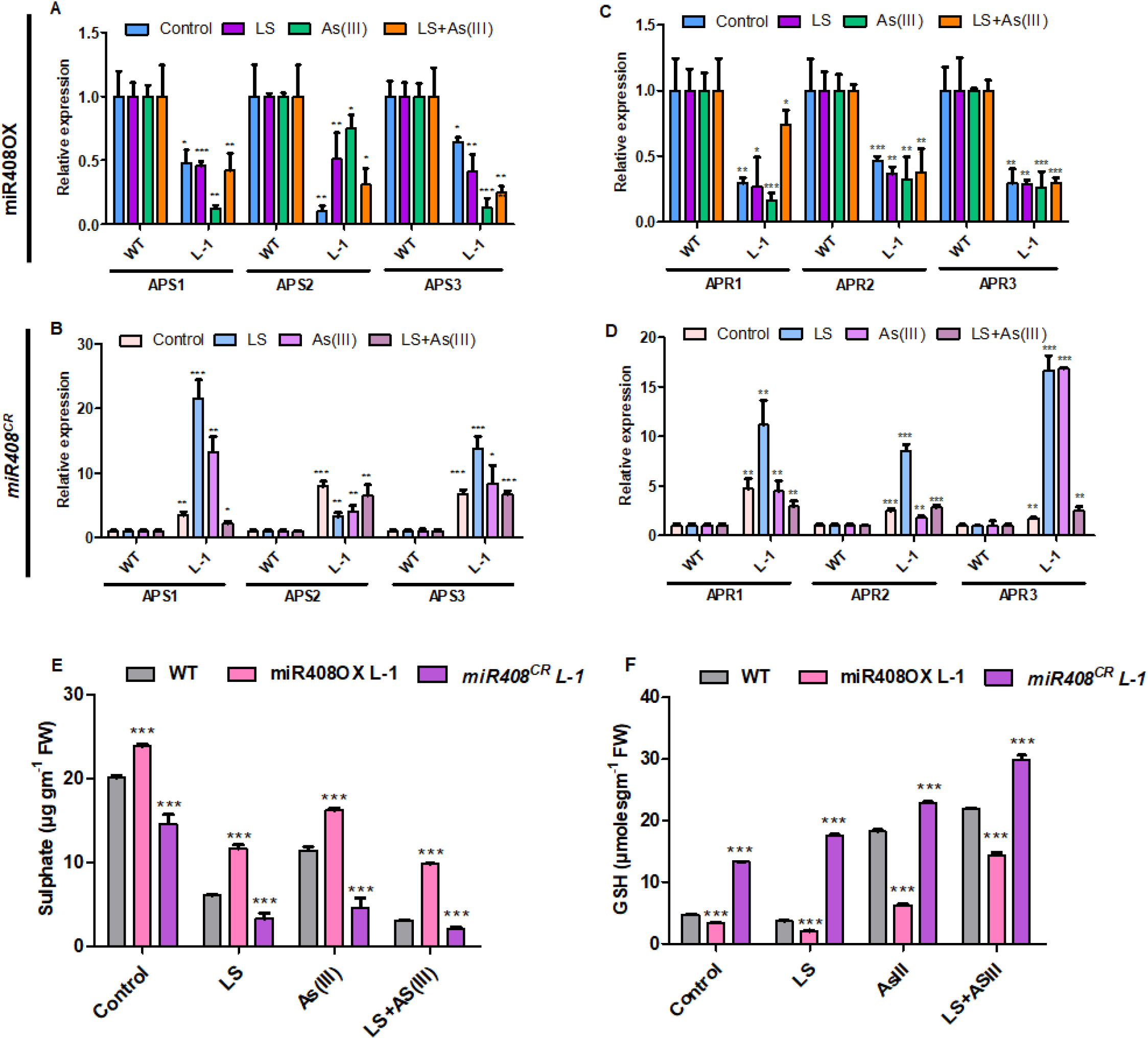
Regulation of sulphur reduction pathway in miR408-OX and *miR408^CR^* plants compare to WT seedlings. **(A)** Relative expression of *AtAPS1*, *2* and *3* in miR408-OX plants under Control, LS, As(III) (10µM), [LS+As(III)] (10 µM). (**B)** Relative expression of *AtAPR1*, *2* and *3* in miR408-OX plants under Control, LS, As(III) (10µM), [LS+As(III)] (10 µM) conditions. (**C)** Expression analysis of *AtAPS1*,2 and *3* in *miR408^CR^* plants under Control, LS, As(III) (10µM), [LS+As(III)] (10 µM). (**D)** Expression analysis of *AtAPR1*, *2* and *3* in *miR408^CR^* plants under Control, LS, As(III) (10µM), [LS+As(III)] (10 µM). (**E**) Total sulphur content of miR408-OX (L-1) and *miR408^CR^* (L-1) ten-days old seedlings under Control, LS, As(III) (10µM), [LS+As(III)] (10 µM) conditions. (F) GSH level of miR408-OX (L-1) and *miR408^CR^* (L-1) ten-days old seedlings under Control, LS, As(III) (10µM), [LS+As(III)] (10 µM) conditions **(F)** GSH level of miR408-OX (L-1) and *miR408^CR^* (L-1) ten-days old seedlings under Control, LS, As(III) (10µM), [LS+As(III)] (10 µM) conditions Data are mean ±SD calculated from three biological replicated. *** represent show significant from control at P < 0.001, according to student’s t-test.

Sulphur enters into the plant roots in the form of inorganic sulphate via sulphate transporters (SULTRs). However, all the four families of SULTRs plays an important role in transportation and translocation of S, only *SULTR1* and *SULTR4* transporters are significantly induced under sulphate deficiency in plants (Kumar et al., 2011). Therefore, to study the function of miR408 in regulating sulphate transportation, we analysed the expression of SULTR in miR408OX and *miR408^CR^* plants. *SULTR1;1* and *SULTR4;1* was identified to be significantly downregulated in miR408OX plants under control and stress conditions (Supplemental Figure S9A and S9B), but mutated plants showed significantly increased accumulation **(Supplemental S9C and S9D)**. These results suggest that there must be a higher accumulation of sulphur in the mi408OX plant cells compared to the mutated plant in control and stress conditions.

From the above results, it was clear that the miR408OX plants and *miR408^CR^* plants have differential regulation of S assimilation pathway genes. So, it was very important to measure the initial and the end product of this pathway, which are sulphur and GSH. Interestingly, higher sulphur accumulation was found in the miR408OX plant compared to the WT in LS, As(III) and [LS+As(III)] stress conditions. However, sulphate content in *miR408^CR^* plants was lower as compared to WT (Figure 5E, Supplemental Figure S10A and S10B). This result clearly suggests that mutated plants have better sulphate assimilation than miR408OX plants and therefore are more tolerant to low sulphur and As stress.

We estimated Glutathione (GSH), which acts as damage control after stress-induced oxidative damage in plants (Kumar et al., 2018; Hasanuzzaman et al., 2019). GSH content was lower in miR408OX plants which clearly indicates that they are more susceptible to stress conditions and are more sensitive. At the same time, increased GSH was observed in *miR408^CR^* plants in control as well as stress condition, which clearly indicates that mutated plants have a better detoxification mechanism than miR408OX plants (Figure 5F, Supplemental Figure S11A and S11B).

### miR408 encodes functional peptide miPEP408-35aa

We analysed upstream from the mature miR408 sequence to identify putative ORFs encoding miPEP408. Analysis revealed presence of two putative ORFs (129 bp and 108 bp) (Supplemental Figure S12). These ORFs are designated as ORF_1_ and ORF_2_ based on their location with respect to the mature miR408. The ORF immediately upstream of miR408 was designated ATG1 and the next ORF was called ATG2. These ORFs may encode small peptides of 42 aa and 35 aa, respectively, designated as miPEP408-42aa and miPEP408-35aa (Figure 6A). To know the functional ORF, seedlings were grown on media supplemented with exogenous miPEP408-42aa and miPEP408-35aa at different concentrations (0.25 to 1 μM), as it has been well established that the exogenous application of miPEPs enhanced the transcript of pri-miRNA and associated phenotype. The miPEP408-35aa showed a concentration-dependent increase of seedling growth and the expression of pri-miR408, while the miPEP408-42aa supplemented seedlings did not show a significant change in the growth of the seedlings and the expression of pri-miR408 compared to the control condition (Figure 6B-6E). The transcript levels of miR408 targets, like *Plantacyanin (ARPN)* and *Laccase3 (LAC3)* were downregulated in seedlings grown on miPEP408-35aa supplemented media compare to control whereas miPEP408-42aa did not affect the modulation in the expression of targets genes (Figure 6F-6G). These results suggested that miPEP408-33aa might be the functional miPEP encoded by a pri-miR408 have the potential to enhance the expression of miR408 and modulate associated phenotype. Preliminary dose-dependent experiments represented that 0.50 μM concentration of miPEP408-35aa was optimal and selected for further studies of miPEP408-mediated regulation of miR408.

**Figure 6.**
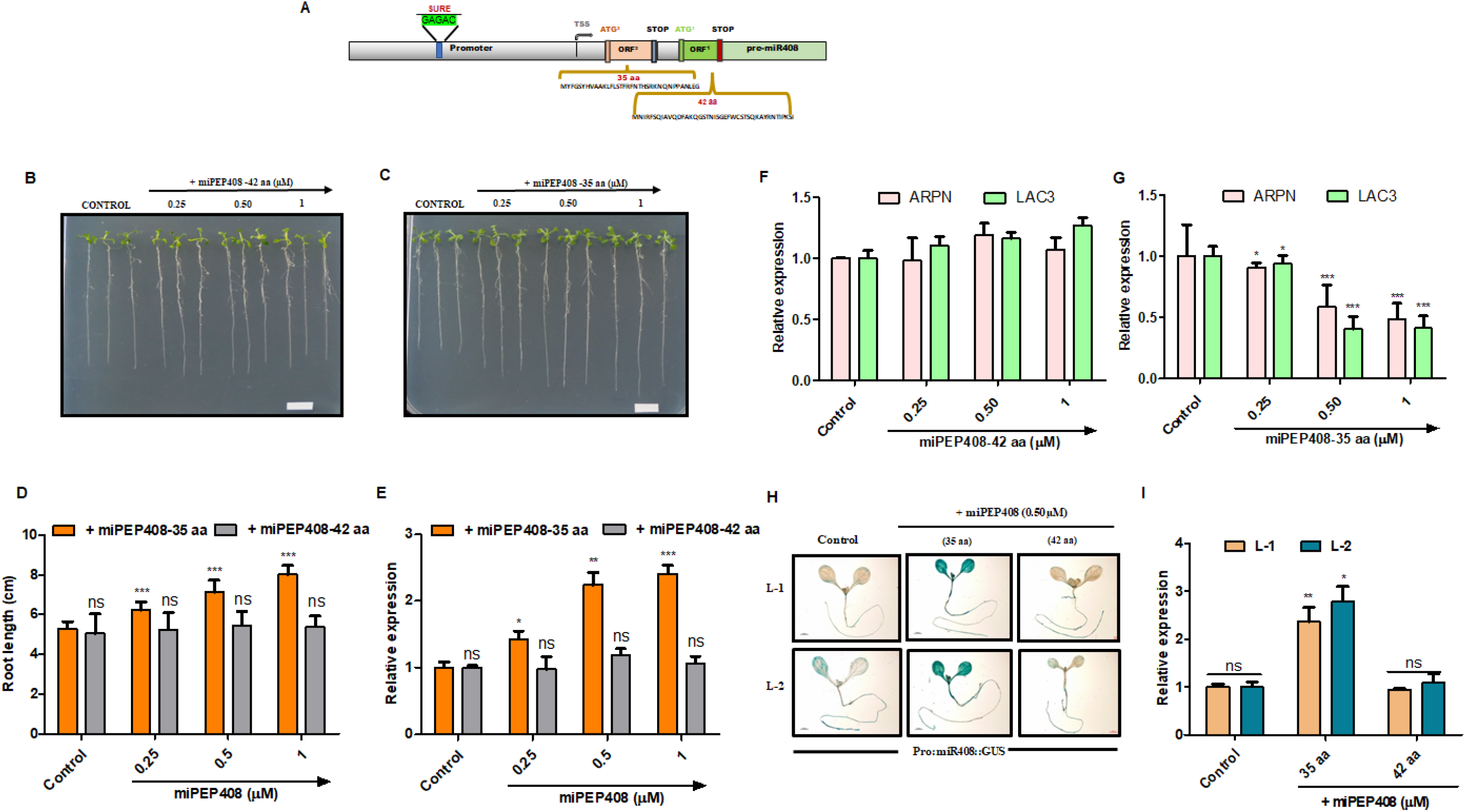
Identification and functional characterization of miPEP408. (**A)** Schematic representation of SUREs element and different ORF found on the promoter of pri-miR408 with predicted TSS and precursor sequence. (**B)** Effect of ten-day-old WT seedlings grown on half-strength MS medium supplemented with water (control) or different concentrations of miPEP408-42 aa. Scale bar, 1 cm. (**C)** Effect of ten-day-old WT seedlings grown on half-strength MS medium supplemented with water (control) or different concentrations of the miPEP408-35 aa Scale bar, 1 cm. (**D)** Measurement of root lengths of ten-day-old WT seedlings grown on half-strength MS medium supplemented with water (control) or different concentrations of miPEP408 (35 aa & 42 aa). n = 30 independent seedlings. (**E)** Expression analysis of pre-miR408 in ten-day-old WT seedlings grown on half-strength MS medium supplemented with water (control) and different miPEP408 (35 aa & 42 aa) concentrations. (**F, G)** Expression analysis of ARPN and LAC3 in ten-day-old WT seedlings grown on half-strength MS medium supplemented with water (control) and different miPEP408 (35 aa & 42 aa) concentrations. (**H)** Histochemical GUS staining of five-day-old promoter line seedlings of Pro:miR408::GUS grown on half-strength MS medium supplemented with water (control), miPEP408-35 aa and miR408-42 aa (0.50μM). Scale bars, 1,000μm. **(I)** Expression of GUS transcript in five-day-old seedlings of Pro:miR408::GUS grown on half-strength MS medium supplemented with water (control), miPEP408-35 aa and miR408-42 aa (0.50μM). Calculations of data were performed from three biological replicated independently per treatment with similar results. The asterisk denotes a significant difference in values with *** as P<0.001, according to the two-tailed student’s t-test.

To provide better information of functional miPEP further, promoter-reporter lines (*PromiR408::GUS*) were developed to understand the role of miPEP to the responsiveness of the miR408 promoter. The promoter lines were grown on a medium supplemented with or without synthetic miPEP408-42aa and miPEP408-35aa. Histochemical GUS assay revealed enhanced GUS activity in *PromiR408::GUS* lines supplemented with miPEP408-35 aa, not with miPEP408-42aa, compared to control (Figure 6H). Similarly, increased expression of the GUS gene was only observed in the seedlings grown in media supplemented miPEP408-35aa, not with miPEP408-42aa, compared to control (Figure 6I). These results collectively suggest that the pri-miR408 encodes a small functional peptide of 35 amino acids.

### Application of miPEP408 leads to growth inhibition and exhibits miR408OX-associated phenotypes under stress conditions

From the above experiments, it is understandable that miPEP408 transcriptionally regulates the miR408 expression, however, the impact of miPEP on different stress was still not studied. Hence, we added exogenous miPEP408-35aa peptide (0.50 µm) in control and LS, As(III) and [LS+As(III)] media. Intriguingly, the supplementation of the exogenous peptide to the ½ MS media increased the sensitivity of seedlings towards LS, As(III) and [LS+As(III)] stress, which was evident from the greater reduction in primary root length and fresh weight (Figure 7A-7C). This result indicates that the addition of miPEP408 negatively regulates the plant growth in LS, As(III) and [LS+As(III)] stress in Arabidopsis and behave similarly to miR408OX plants.

**Figure 7.**
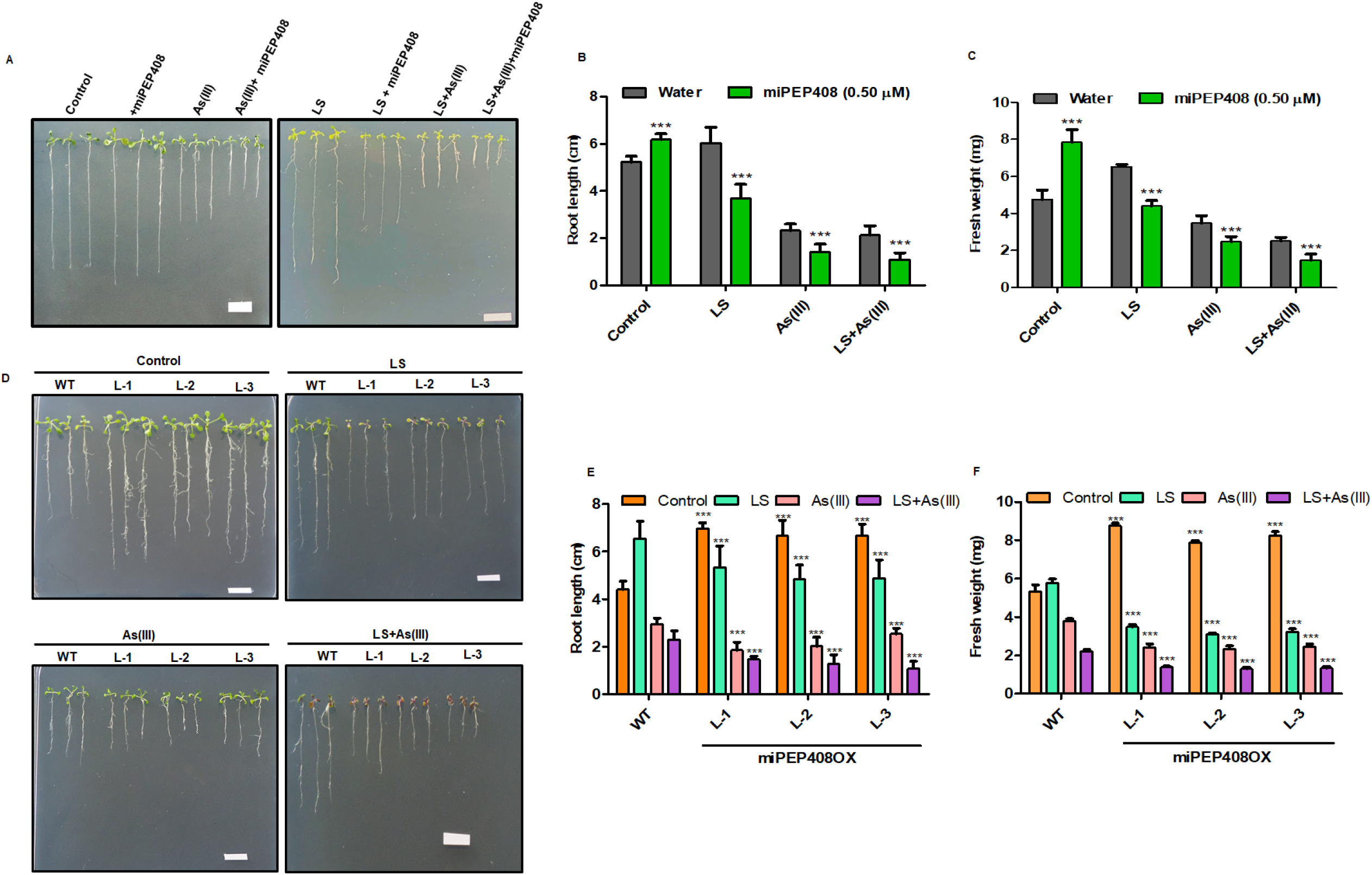
Exogenous application and overexpression lines of miPEP408 exhibit miR408-OX associated phenotype in stress conditions. **(A)** Representative image of ten-day-old WT *Arabidopsis* seedlings grown on control, LS, As(III) and [LS+As(III)] conditions supplemented with water and miPEP408-35 aa (0.50µM) separately in each condition. Scale bar, 1 cm. **(B**) Primary Root lengths of ten-day-old WT seedlings grown on control, LS, As(III) and [LS+As(III)] conditions supplemented with water and miPEP408-35 aa (0.50µM) separately in each condition. N=30 independent seedlings. (**C)** Fresh weight of ten-day-old WT seedlings grown on control, LS, As(III) and [LS+As(III)] conditions supplemented with water and miPEP408-35 aa (0.50µM) separately in each condition. N=30 independent seedlings. **(D)** Representative image of ten-day-old WT and miPEP-OX seedlings grown on control, LS, As(III) and [LS+As(III)] conditions. Scale bar, 1 cm. **(E)** Primary Root lengths of ten-day-old WT and miPEP408-OX seedlings grown on control, LS, As(III) and [LS+As(III)] conditions. N=30 independent seedlings. **(F)** Fresh weight of ten-day-old WT and miPEP408-OX seedlings grown on control, LS, As(III) and [LS+As(III)] conditions. N=30 independent seedlings. This data was generated from three biological replicated independently per treatment with similar results. The asterisk denotes significant difference in values with *** as *P < 0.1; **P < 0.01; ***P < 0.001, according to two-tailed student’s t-test.

To briefly elucidate the miPEP408 function in stress conditions, we developed overexpressing miPEP408 overexpressing lines. These lines accumulated miR408 and decreased the accumulation of its target *ARPN* expression (Supplemental Figure S13A and 13B). To functionally validate the role of miPEP408 in LS and As stress, miPEP408OX lines were grown on LS, As(III) and combined [LS+As(III)] conditions. Phenotypic analysis revealed that miPEP408OX plants show better growth represented as root length and fresh weight compared to the WT whereas seedlings grown under LS, As(III) and [LS+As(III)] conditions showed growth inhibition as a decreased primary root length and fresh weight compared to WT (Figure7D-7F). These results clearly show that exogenous miPEP408 and miPEP408OX plants exhibit similar phenotypes as miR408OX plants in control and stress conditions (Figure 3E-3G).

### Differential ROS, sulphur accumulations and higher lipid peroxidation exhibit exogenous miPEP408 and miPEP408OX plants

To further study the effect of miPEP408 on the antioxidant machinery, miPEP408OX plants and the WT seedlings were grown on media supplemented with miPEP408 analyzed for ROS accumulation under different stresses. NBT staining results suggested a higher accumulation of O^-^ radical in seedlings grown with exogenous peptide and miR408OX in LS, As(III) and [LS+As(III)] stress. Additionally, no significant NBT staining was seen in seedlings grown on control media (supplemented with either water or peptide) and miPEP408OX under control conditions (Figure 8A and 8B). In stress conditions, there was a higher accumulation of free radicals in seedlings supplemented with the miPEP408, which behaved like the miR408OX plants (Figure 4A). Furthermore, DAB staining showed that there was higher brown precipitate in the peptide supplemented and miPEP408OX seedlings under LS, As(III) and LS+As(III) stress compared to the control (Supplemental Figure S14A and S14B). These results suggest that seedlings grown on peptide supplemented media have a higher accumulation of ROS and are more sensitive to the stress compared to the seedlings grown under stress conditions with control (water) as a supplement. From these results we concluded that miPEP408 supplemented seedlings are behaving like miR408OX plants under LS, As(III), [LS+As(III)] stress.

**Figure 8.**
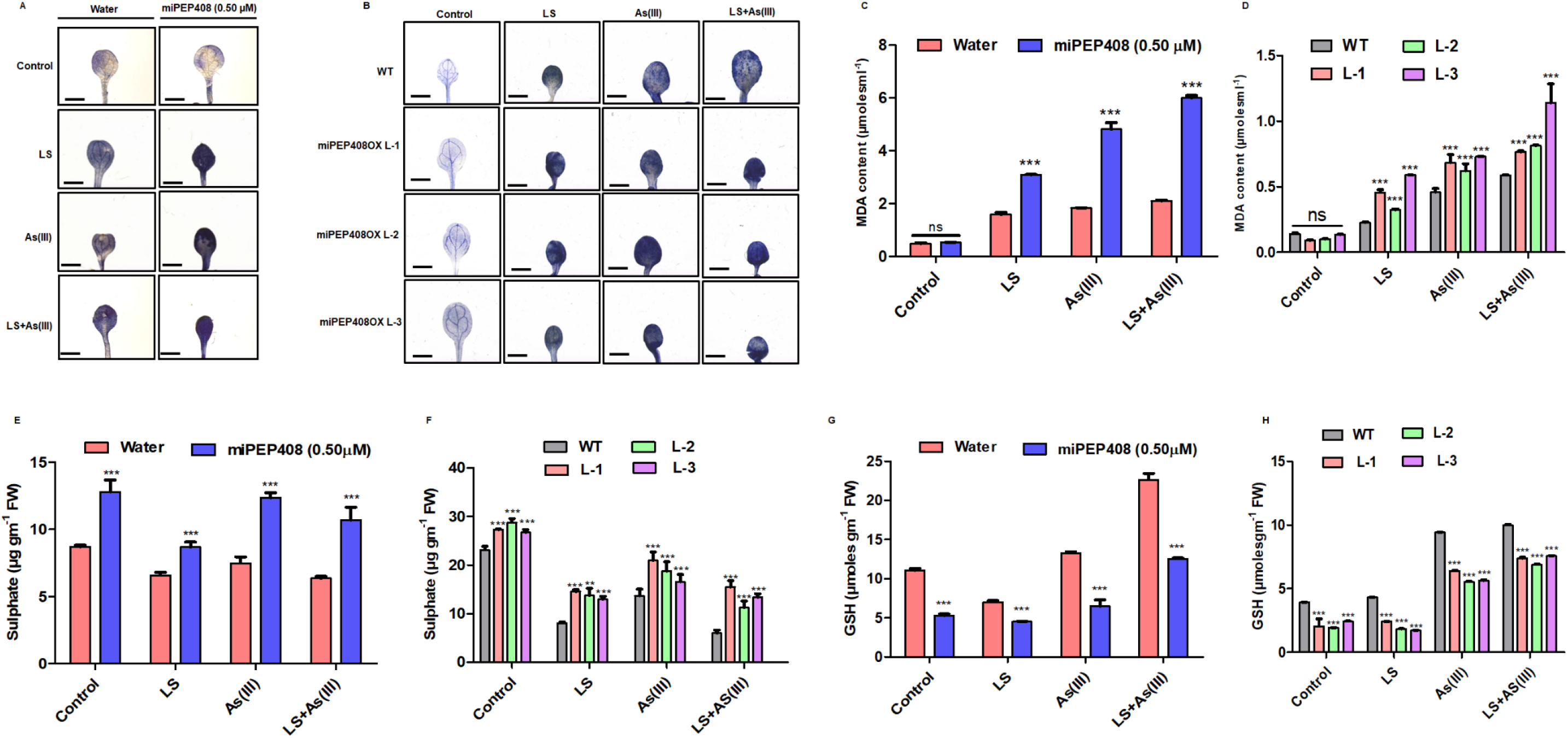
Exogenous miPEP408 and miPEP408-OX show differential ROS accumulation Sulphur level and lipid peroxidation in plants in limiting sulphur and As(III) stress. Ten-day-old WT, miR408-OX and miR408CR were grown for 10 days on medium containing optimum sulphur as control (C), limiting sulphur (LS), As (III) (10 μM), [LS + As (III)] (10μM). **(A)** Nitrotetrazolium (NBT) staining of WT *Arabidopsis* seedlings grown for 10 days on control and stress conditions supplemented with Water and miPEP408 separately. **(B)** Staining of WT and miPEP408-OX seedlings with Nitrotetrazolium (NBT) in control and stress conditions. **(C)** Total MDA content of Ten-day-old WT seedlings was grown for 10 days on control and stress medium containing supplemented with Water and miPEP408 separately. **(D)** Estimation of MDA in WT, miPEP408-OX seedlings in control and stress conditions. **(E)** Total sulphur content ten days grown WT seedlings grown under control and stress conditions LS, supplemented water and miPEP408 individually. **(F)** The total sulphur content of miPEP408-OX ten-days old seedlings under control and stress conditions (**G)** Total GSH content of Ten-day-old WT seedlings grown for ten-days on control and stress medium supplemented with Water and miPEP408 individually **(H)** GSH level of miPEP408-OX ten-days old seedlings under control and stress conditions. This data was generated from three biological replicated independently per treatment with similar results. The asterisk denotes significant difference in values with *** as *P < 0.1; **P < 0.01; ***P < 0.001, according to two-tailed student’s t-test. bar=1mm.

MDA content was also estimated in with or without peptide supplemented and miPEP408OX seedlings under LS, As(III), [LS+As(III)] stress. Higher accumulation of MDA was observed in the miPEP408OX and seedlings grown on miPEP408-35aa supplemented media along with LS, As(III) and [LS+As(III)] stress compared to the water added growth media (Figure 8C and 8D). There was no significant change in the control condition. These results suggest that miPEP408 enhances the sensitivity compared to the control and shows a similar phenotypic effect as miR408OX plant on LS, As(III) and combined [LS+As(III)] stresses. To, observe the sulphur availability inside the cell, we measured the sulphur content of 10-day-old different lines of miPEP408OX and WT seedlings grown on control and peptide supplemented media, under LS, As(III) and [LS+As(III)] stress conditions. Results imply that miPEP408OX plants and exogenous peptide treatment enhances the sulphate content in control and stress conditions in Arabidopsis (Figure 8E and 8F). Measurement of GSH content in the miPEP408OX lines and peptides supplemented seedlings suggest that the exogenous application of miPEP408 and miPEP408OX lines have decreased GSH content in control and stress conditions (Figure 8G and 8H).

### miPEP408 regulates sulphur metabolism and affects plant tolerance to stress

The above results clearly suggested that miR408 negatively regulates the sulphate assimilation pathway. Therefore, to know whether exogenous application of miPEP408 also impacts sulphate reduction, seedlings grown on media having LS, As(III) and [LS+As(III)] stress along with miPEP408 were used for expression analysis of sulphur assimilation related genes (*APS and APR*). The accumulation of *AtAPS1*, *AtAPS2*, and *AtAPS3* is negatively affected in all the stress conditions (Supplemental Figure S15A). This result shows that sulphur is not being used in the reduction pathway and eventually not be used for the synthesis of GSH needed for providing tolerance to plants. The expression of *AtAPR1,2* and *3* was also significantly decreased in the peptide supplemented seedlings in control and stress conditions (Supplemental Figure S15B). Analysis of APS and APR genes in control conditions in miPEP408OX lines showed that accumulation of miPEP408 also affects the sulphur assimilation pathway genes and affect sulphur assimilation (Supplemental Figure S16A and S16B). From these results now it is confirmed that miPEP408 regulates the sulphur assimilation pathway gene and might affect the stress tolerance mechanism as shown by miR408 overexpressing plants previously.

This result suggests that miPEP408 either regulates S uptake or its transportation in the cell. As mentioned in previous results, apart from S assimilation genes, sulphate transportation mediated by SULTRs also strictly regulates plants’ responses towards S limitation and As stress. Therefore, the expression of SULTR1 and SULTR4 was analysed in presence of miPEP408 in Arabidopsis. Interestingly, expression profiling revealed a decrease in transcripts of the isoforms associated with both the SULTRs on the application of the exogenous miPEP408 under LS, As(III) and [LS+As(III)] stress (Supplemental Figure S15C). These results showed that the application of the miPEP408 increases sensitivity in plants under LS and As stress conditions.

### miR408 modulates the arsenic accumulation

Various reports suggest As toxicity is alleviated by sulphur by affecting the accumulations by modulating amino acids and thiol metabolism in rice (Dixit et al., 2015). Therefore, we observed the effect of miR408 on As accumulation in miR408OX and *miR408^CR^* plants and those supplemented with miPEP408 under control, LS, As(III) and LS+As(III) conditions. It was observed that miR408OX plants have a significantly lower accumulation of total As under As(III) and [LS+As(III)] compared to *miR408^CR^* plants (Supplemental Figure S17A and 17B). The miPEP408 supplemented seedlings grown on As(III) and LS+As(III) also showed a higher accumulation of As, similar to miR408OX plants (Supplemental Figure S17C).

## Discussion

MicroRNAs are small RNAs functioning in the regulation of all the processes of plants that include development and responses to abiotic stresses by posttranscriptional inhibition of target transcript (Vakilian et al., 2020; Millar et al., 2020; Pagano et al., 2021). Recent studies showed that the small peptide encoded by some of the pri-miRNAs participate in the regulation of miRNA expression (Lauressergues et al., 2015; Couzigou et al., 2015; Sharma et al., 2020; Prasad et al., 2020; Badola et al., 2022). However, detailed insight into the role of these peptides under different stresses is still lacking. Among the known families of miRNA, miR408 is identified to be conserved and has a fundamental function with its diverse response to nutrient limitation, cold, drought, osmotic and oxidative stress (Ma et al., 2015; Song et al., 2018). This study demonstrated that miR408-edited plants showed a higher degree of survival than miR408 overexpressing plants in nutrient deficiency and As stress. Moreover, the function of the peptides encoded by pri-miR408 was studied in detail and the study suggests that miPEP408 regulates tolerance towards abiotic stresses such as sulphate deficiency and As toxicity.

Small RNA sequence analysis concluded that 55 miRNAs are differentially regulated in As and LS conditions, out of which only 18 miRNAs were common in these stresses (Figure 1 **and** Supplemental Table S5 and S6). Of common differentially expressed miRNAs, miR408 was the most responsive miRNA towards As and LS conditions (Supplemental Figure 1). The expression of miR408 is differentially regulated in various plant systems in response to different abiotic stresses, like drought, As and nutrient deficiencies (Sharma et al., 2015; Vasupalli et al., 2020; Liang et al., 2015). We analysed the involvement of miR408 towards LS, As(III), [LS+As(III)] stress and observed significant downregulation in the expression of miR408 and vice-versa of its targets, ARPN and LAC3 (Figure 1C and 1D). Since sulphate limitation and As toxicity were identified to regulate the expression of miR408, the activity of the promoter of the miR408 was studied under these stresses. Previous studies have reported that environmental constraints, including heavy metal and nutrient alteration, tightly regulate the promoter activity of responsive gene regulation, thereby affecting plant growth and development (Qi et al., 2007; Tiwari et al., 2020). Expectedly, reduced transcript levels and GUS activity in *PromiR408::GUS* stable lines under stress conditions compared to EV were observed (Figure 2A-B). Along with the promoter activity, miR408OX lines and CRISPR-edited plants (*miR408^CR^*) of miR408 were developed to have a detailed understanding of the role of this miRNA in stress conditions. Captivatingly, miR408OX lines showed sensitivity towards LS, As(III) and combined [LS+As(III)] stress compared to the WT plants (Figure 3E-3G). Response of miR408OX lines was consistent with previous studies showing reduced growth in plants overexpressing miR408 under drought and osmotic stress in *Arabidopsis* (Ma et al., 2015). Although the study by (Ma et al., 2015) revealed improved growth of plants overexpressing miR408 under salinity, cold and oxidative stress, the response in our study might be specific to As stress and LS. On the other hand, miR408*^CR^* plants were observed to be tolerant towards sulphate limitation and As stress as the root length and fresh weight of mutated plants was significantly increased under LS, As(III) and [LS+As(III)] stress compared to the WT (Figure 3H-3J). Previously CRISPR-edited plants of miR408 were developed in rice however their response under different abiotic stresses was not defined (Zhou et al., 2017). Abiotic stresses such as nutrient deficiency and heavy metal stress increases production of reactive oxygen species (ROS) in plants (Hasanuzzaman et al., 2020). ROS, including superoxide radical (O^2-^), hydrogen peroxide (H_2_O_2_), hydroxyl radical (OH), etc., acts as signal transducers in plants, however, excess levels lead to membrane damage (also termed as lipid peroxidation) and disrupt nucleic acids including DNA and RNA (Choudhury et al., 2017; Singh et al., 2019; Mittler et al., 2017). Increased NBT and DAB staining was observed in miR408OX lines which suggests a higher ROS production in the overexpression plants compared to WT (Figure 4A-4B). This might be a plausible reason for the sensitivity of miR408OX under stress conditions. Additionally, *miR408^CR^* plants accumulated lesser of ROS compared to the WT, which gave clear evidence that miR408 negatively regulates tolerance towards sulphate deficiency and As stress (Figure 4A-4B). A significantly enhanced level of MDA was observed in miR408OX lines compared to the WT under LS, As(III) and [LS+As(III)] conditions. *miR408^CR^* plants accumulated lesser MDA compared to the WT (Figure 4C). Decreased ROS accumulation in miR408*^CR^* plants determines enhanced tolerance against the LS, As(III) and combine [LS+As(III)] condition.

Previous studies revealed that an increase in sulphur reduces As toxicity and also improves thiol metabolism in plants (Dixit et al., 2015). miR408OX and *miR408^CR^* plants showed differential expression of APS and APR genes compared to the WT (Figure 5A-D). The miR408OX plants showed significantly decreased expression of *APS*, *APR* isoforms clearly indicating that the sulphur reduction pathway was negatively regulated in the overexpression plants with reduced accumulation of sulphur and GSH compared to the WT. Interestingly, the mutated plants showed an increased accumulation of various isoforms of *APS* and *APR* transcripts, which clearly showed that there must be a higher accumulation of GSH and more reduction of sulphate, which will provide tolerance to the plant against As stress and LS. Glutathione (GSH) is a sulphur-containing compound having redox and nucleophilic properties that are regulated by sulphur availability and plays an important role in minimizing the stress-induced oxidative damage in plants (Mendoza-Cózatl et al., 2006; Banerjee et al., 2019; Pei et al., 2019). The estimation of sulphur and GSH in miR408OX and *miR408^CR^* plants clearly indicated that the sulphur reduction pathway is regulated by the miR408 expression. Overexpression of miR408 led to a higher accumulation of sulphur and lower accumulation of GSH, indicating that the sulphur reduction pathway is negatively regulated by miR408 might be one of the reasons for sensitive phenotype against stress. *miR408^CR^* plants had greater sulphur reduction (lesser sulphate content) and high GSH content suggesting that these plants have highly efficient sulphur reduction metabolism capability and better use of the sulphur for the GSH biosynthesis compared to the WT (Figure 5E and 5F). This result is in corroboration with previous studies in which tolerant natural accession was identified to have lesser sulphate and greater GSH content under LS, As(III) and combine [LS+As(III)] stress (Khare et al., 2017). These analyses suggest that the absence of miR408 confers tolerance to the *miR408^CR^* plants towards LS and As stress.

To identify ORFs encoded by miR408, sequence upstream was analysed and ORFs encoding putative peptides of 35aa and 42aa were identified in pri-miR408 (Supplemental Figure 12). Interestingly, exogenous application of miPEP408-35aa, but not miPEP408-42aa, increased primary root growth of the WT plants as well as accumulation of miR408 (Figure 6A-6D). The impact of miPEP408-35aa was also reflected in the expression of miR408 and its targets, APRN1 and LAC3, whose expression got reduced after peptide treatment compared to the control (Figure 6F-6G). This result was in accordance with the previous research where overexpression of miR408 in rice led to increased growth and development (Zhang et al., 2017).

Histochemical GUS assay illustrated increased GUS activity along with increased GUS expression in *PromiR408::GUS* lines when grown on media supplemented with miPEP408 35aa compared to control, but not with miPEP408-42aa (Figure 6H-6I). Similar results were observed in the study by Sharma et al., (2020), which showed transcriptional regulation of miR858 via miPEP858a. Various studies suggest that small peptides might play a key role in abiotic stress tolerance like AtPep3 which is a hormone-like peptide that enhances the salinity stress tolerance in Arabidopsis (Nakaminami et al., 2018). Similarly, the C-terminally ENCODED PEPTIDE 5(CEP5) promotes tolerance in Arabidopsis towards osmotic and drought stress by regulating AUX/IAA Equilibrium (Smith et al., 2020). Recently, a few miPEPs have been identified in plants (Lauressergues et al., 2015; Waterhouse et al., 2015) but their role in abiotic stress tolerance is yet to be explored. We demonstrated that the application of miPEP408 on seedlings and miPEP408OX plants grown on the 1/2 MS media grow better in the control condition. Interestingly, when these seedlings were grown on the LS, As(III) and [LS+As(III)] supplemented with water as control or peptide and miPEP408OX lines exhibited contrasting phenotypes. The seedling grown on stress media supplemented with the exogenous miPEP408 as well as miR408OX showed a sensitive phenotype compared to the water supplemented media (Figure 6A-F). Treatment with the exogenous miPEP408 peptide and miPEP408OX seedlings showed increased sensitivity towards heavy metal and nutrient deficiency which was also reflected by the increased NBT and DAB staining along with elevated MDA content in the WT plants supplemented with the exogenous peptide and miR408OX plants under different stresses (Figure 8A-8C).

Exogenous supplementation of the miPEP408 or overexpressing miPEP408 negatively affects the expression of important sulphur reduction pathway genes, like APS and APR, that are required for the synthesis of GSH which is involved in the detoxification mechanism (Supplemental Figure 15 and 16). Reduced sulphur and GSH content were observed in overexpressing lines and peptide supplemented seedlings in control and stress conditions (Fig. 9E and 9H). Analysis suggests that miPEP408 enhance the sensitivity response of *Arabidopsis in* comparison to the normal conditions, thus reducing the tolerant phenotype in stress conditions. Sulphur assimilation is largely regulated by sulphate transporters (SULTRs) in plants. Expression analysis of *SULTR1;1* and *SULTR4;1* revealed that both the genes were downregulated after supplantation of miPEP408 or in miPEP408OX lines (Supplemental Figure 15C). This suggests that miPEP408 supplemented seedlings possess a higher accumulation of sulphur, thereby restricting the uptake of sulphur by SULTRs under LS, As(III) and [LS+As(III)] conditions. Similar results have been also reported in contrasting Arabidopsis natural variants under LS and As stress (Khare et al., 2017). Moreover, a recent study elucidated the importance of SULTR1;1 in providing tolerance against sulphate limitation and As toxicity (Kumar et al., 2019). The As accumulation data suggests that miR408OX and exogenous miPEP408 treated seedlings show lower accumulation of As due to a lower GSH pool and the mutated lines have a higher GSH pool and higher accumulation of As (Supplemental Figure 17). Therefore, we came to the conclusion that miR408 and its associated peptide regulate sulphur metabolism and modulate and reduce glutathione levels to regulate the arsenic detoxification mechanism.

**Figure 9.**
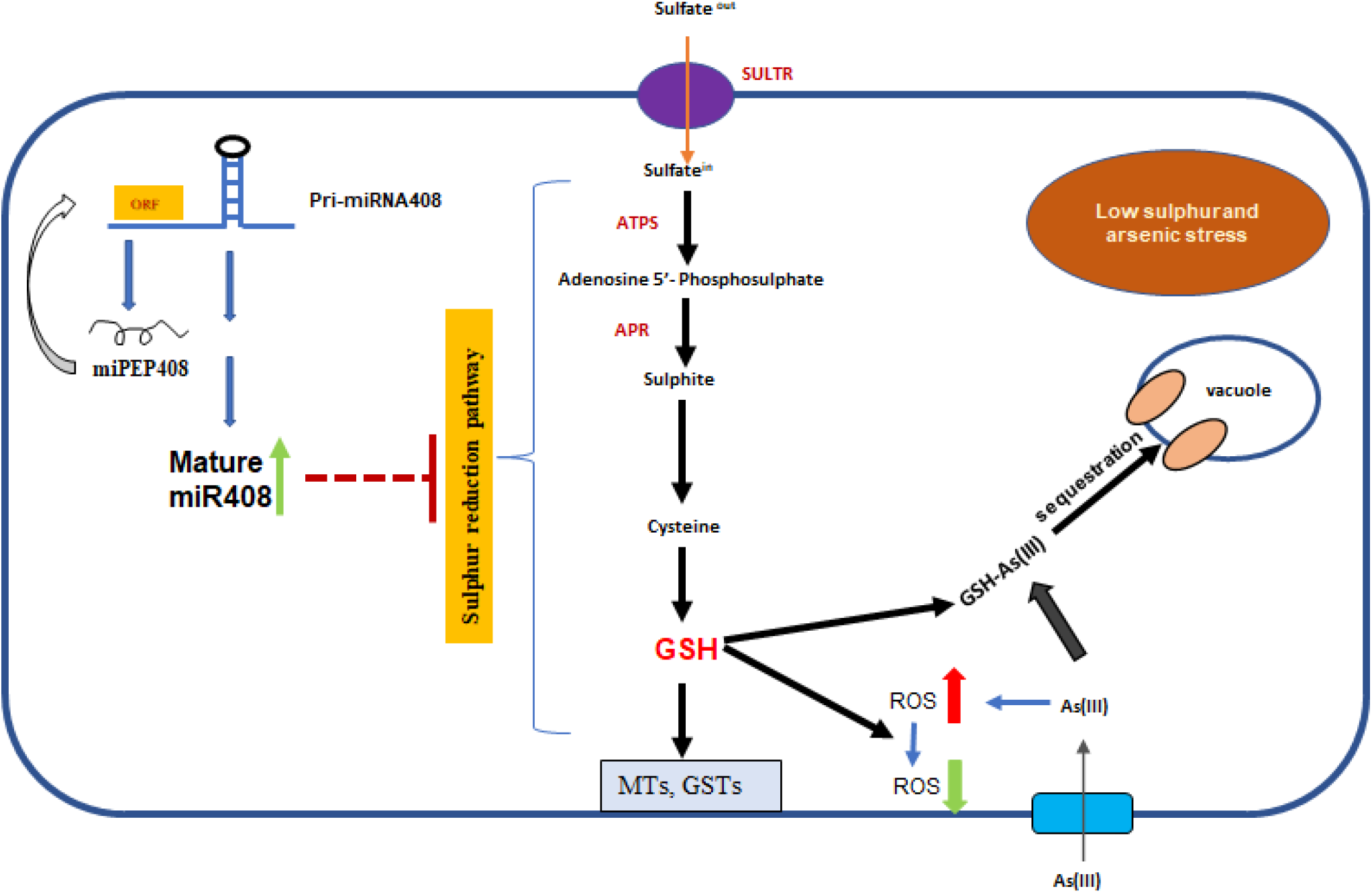
A schematic model representing miPEP408 controls miR408 transcription to mediated regulation of sulphur metabolism under limiting sulphur (LS) and As(III) stress. This model represents the key role of miPEP408 regulating the sulphur reduction pathway via pre-miR408 and mediating the regulation of genes involved in sulphur metabolism. Which clears the regulatory role of miPEP408 in nutrient deficiency and arsenic tolerance. Genes which modulated in the pathway are marked as red.

On the basis of our results, we propose a model representing miPEP408 regulating the expression of miR408, which negatively regulates the sulphur reduction pathway in Arabidopsis under nutrient and As stress leading to its involvement in the detoxification mechanism against heavy metal and nutrient deficiency (Figure 9).

## Method and material

### Plant Materials and Growth Conditions

Plant material and growth conditions Arabidopsis *A. thaliana* (Col-0) was used as the WT plant and for overexpression and the editing of miiR408 through the CRISPR–Cas9 approach. Seeds were surface sterilized and grown on one-half-strength MS medium (Sigma-Aldrich, USA) according to Shukla et al., (2015). For the control condition, an optimum concentration (1500 µM) of sulphate was used in the medium. For limiting sulphate conditions, MgSO_4_ was replaced by MgCl_2_ and a 10 µM concentration of sulphate was supplemented. For As(III) treatment, 10 µM Na_2_AsO_3_ (Stock solution 50 mM Na_2_AsO_3_, ICN, USA) was added to the one-half-strength MS medium (Sigma-Aldrich, USA). The pH of all the media was adjusted to 5.75-5.85 with 0.1 M KOH or HCl. Seeds were grown in a growth chamber (Conviron, USA) under controlled conditions of 16-h-light/8-h-dark photoperiod cycle, 22°C temperature,150 to 180 µmol m^-2^ s^-1^ light intensity, and 60% relative humidity for 10 days.

### Synthetic peptide assay

The synthetic peptides (purity > 95%) were synthesized through Link Biotech (http://www.linkbiotech.com). The peptides were dissolved in water (stock concentration, 5 mM). The seedlings were treated with concentrations from 0.25 to 1 μM peptide diluted in the agar medium or in water.

### Peptide sequences

**miPEP408-42 aa-MNIRFSQIAVQDFAKQGSTNISGEFWCSTSQKAYRNTIPKSI**

**miPEP408-35 aa-MYFGSYHVAAKLFLSTFRFNTHSRKNQNPPANLEG**

### Biochemical analysis

Lipid peroxidation assay Lipid peroxidation is a biochemical marker for oxidative stress, which is estimated by malondialdehyde (MDA) produced using the thiobarbituric acid (TBA) method (Baryla et al., 2000). Briefly, 0.1 g of sample was ground into powder using liquid nitrogen and was homogenized in 1 mL 0.1% (w/v) trichloroacetic acid (TCA). The homogenate was centrifuged at 10,000xg for 10 min. The supernatant was mixed with 4 ml of 20% TCA containing 0.5% TBA, heated in a boiling bath (95^0^C) for 15 min and then allowed to cool rapidly in an ice bath. The mixture was centrifuged at 10,000xg for 5 min and the resulting supernatant was used for the determination of MDA content. The concentration of MDA was calculated by measuring absorbance at 532 nm (correction was made by subtracting absorbance at 600 nm for turbidity) by using an extinction coefficient of 155 mM^−1^ cm^−1^.

### Histochemical GUS staining

The GUS staining was performed using a previously described method (Jefferson et al., 1989). Seedlings of the *PromiR408::GUS* transgenic lines were immersed in a solution containing 100 mM sodium phosphate buffer (pH 7.2), 10 mM EDTA, 0.1% Triton X-100, 2 mM potassium ferricyanide, 2 mM potassium ferrocyanide and 1 mg ml^−1^ 5-Bromo-4-chloro-3-indolyl-β-D-glucuronide at 37 °C for 4 h. The chlorophyll was removed by incubation and multiple washes using 70% ethanol. The seedlings were observed under a Leica microscope (LAS version 4.12.0, Leica Microsystems) for the GUS staining.

### Histochemical detection of superoxide and hydrogen peroxide accumulation

NBT staining was used to detect the production of superoxide radicals and was carried out according to the method (Jabs et al.,1996). Ten-day old seedlings were immersed in 50 mM potassium phosphate buffer (pH 7.8) containing 0.1 mg ml^-^1 nitrobluetetrazolium (NBT) at 250C in the dark for 2 h. Stained samples were bleached in 80% ethanol and incubated at 700C for 10-20 min. To analyze the accumulation of hydrogen peroxide (H_2_O_2_) in the samples, DAB staining was performed according to the method (Daudi et al., 2012). Ten-day-old seedlings were immersed in 1 mg ml^-1^ DAB solution. To enhance the uptake of DAB, samples were vacuum infiltrated for 5-10 min. Staining was carried out in dark for 4-5 h with mild shaking. Finally, the samples were de-stained using a 1:1:3 mixture of acetic acid, glycerol and ethanol at 95°C for 15 min. Seedlings were visualized using a Stereoscope zoom binocular microscope (Leica LAS version 4.12.0, Leica Microsystems).

### Sulphur estimation

Plant seedlings (200 mg) were homogenized in 4 ml of 0.1 M HCl for 2 h at room temperature. After centrifugation at 12,000xg supernatant was recovered and used for the determination of sulphate content using the turbidimetric method (Tababai et al., 1970). The standard curve was plotted using sodium sulphate as a standard and the readings were recorded at 420 nm.

### Reduced glutathione (GSH) estimation

A homogenate of ten-day-old seedlings (0.5 g) was prepared in 2.5 ml of 5% TCA. Homogenate was centrifuged at 4,000xg for 10 min. The supernatant (0.1 ml) was used for the estimation of GSH. The supernatant (0.1 ml) was made up to 1.0 ml using 0.2 M sodium phosphate buffer (pH 8.0) followed by the addition of 2.0 ml of freshly prepared DTNB (5,5′-dithiobis (2-nitrobenzoic acid) solution. The concentration of GSH was determined spectrophotometrically at 412 nm after 10 min of incubation (Moron et al., 1979). The values are expressed as µmoles g ^-1^ sample.

### RNA isolation and gene expression analysis

Ten-day old seedlings were used to isolate total RNA using Spectrum Plant Total RNA Kit (Sigma–Aldrich, USA) as per the manufacturer’s instructions. RNA quantity and quality were analysed using NanoDrop spectrophotometer (NanoDrop, Wilmington, DE, USA) and agarose gel electrophoresis. RNase-free-DNase-I (Fermentas, LifeSciences, Canada) was used to remove DNA contamination. For the cDNA preparation, Revert Aid First Strand cDNA synthesis kit (Fermentas, LifeSciences, Canada) was used to reverse transcribe total RNA. Quantitative Real Time-PCR was carried out in an ABI7500 instrument (ABI Biosystems, USA) using SYBR Green Supermix (ABI Biosystems, USA). As an internal control, the tubulin gene was used to quantitate the relative transcript level of the genes of interest using gene-specific oligonucleotides. Data were analyzed using the comparative Ct (2^−ΔΔ^ct) method (Schmittgen et al., 2008).

### Small RNA sequencing and analysis

For the small RNA sequencing, the plant samples were prepared after 10 days of low sulfur and arsenic treatment along with the control condition. The NEBNext Small RNA Sample Preparation protocol is used to prepare a sample sequencing library. The library is prepared as per the kit protocol. The adapters are ligated to each end of the RNA molecule and an RT reaction is used to create single-stranded cDNA. The cDNA is then PCR amplified using a common primer and a primer containing one of 48 index sequences. After libraries are constructed, sequencing was performed on HiSeq 2500 with 1×50bp reads to obtain 40-50 million reads. After the sequencing run, Illumina small RNA-Seq data was processed to generate FASTQ files.

### Statistical analysis

Data are plotted as means ±SD with error bars as standard deviation. The statistical tests and *n* numbers, including sample sizes or biological replications, are described in the figure legends. All the statistical analyses were performed using two-tailed Student’s *t*-tests using GraphPad Prism version 8.4.3 software (* P < 0.1; ** P < 0.01; *** P < 0.001). All the experiments were repeated at least three times independently, with similar results.

## Acknowledgements

We thank Q.-J. Chen at China Agriculture University for the pHSE401.

## Funding

This research was supported by the Council of Scientific and Industrial Research (CSIR), New Delhi, in the form of NCP project no. MLP006. P.K.T. also acknowledges the Department of Biotechnology, New Delhi, for financial support in the form of projects on pathway engineering and genome editing. PKT also acknowledges Department of Science and Technology for the JC Bose. P.K.B., A.S., and H.G. acknowledge the Council of Scientific and Industrial Research, New Delhi, for a Senior Research Fellowship.

## Author contributions

R.S.K. and P.K.T. conceived and designed the study. R.S.K., H.S., and T.D. participated in the execution of all the experiments. R.S.K., H.S. T.D. M.H.A. and P.K.T. interpreted and discussed the data. R.S.K., H.S., T.D., and P.K.T. wrote the manuscript.

## Competing interests

The authors declare that they have no competing interests.

## SUPPLEMENTAL INFORMATION

### Supplemental Data

**Supplemental Figure S1.**
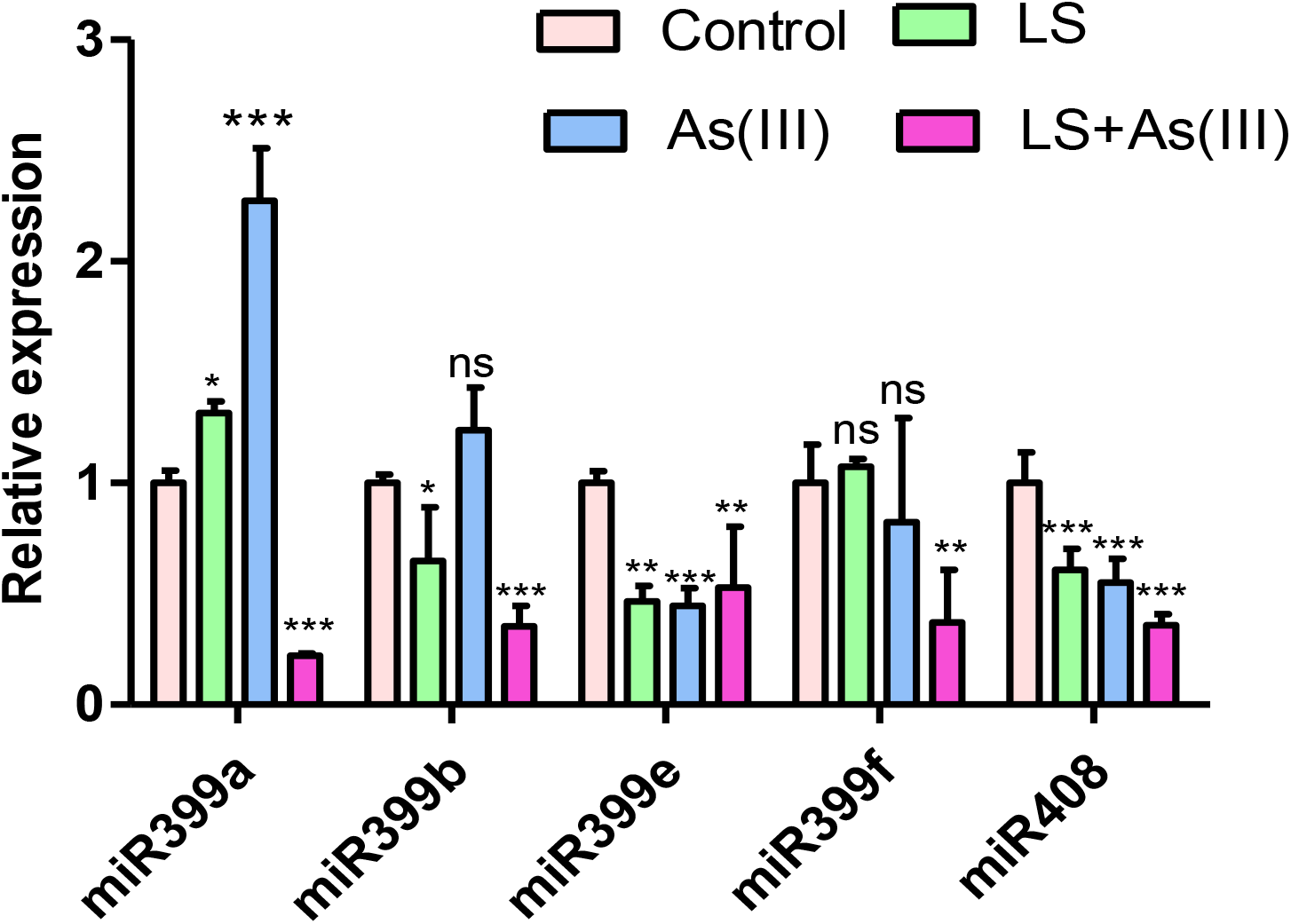
Expression analysis of miRNA(s) under LS, As(III) and LS+As(III) conditions compared to control. Data are mean ±SD calculated from three biological replicated. *** represent significant from control at P < 0.001, according to student’s t-test.

**Supplemental Figure S2.**
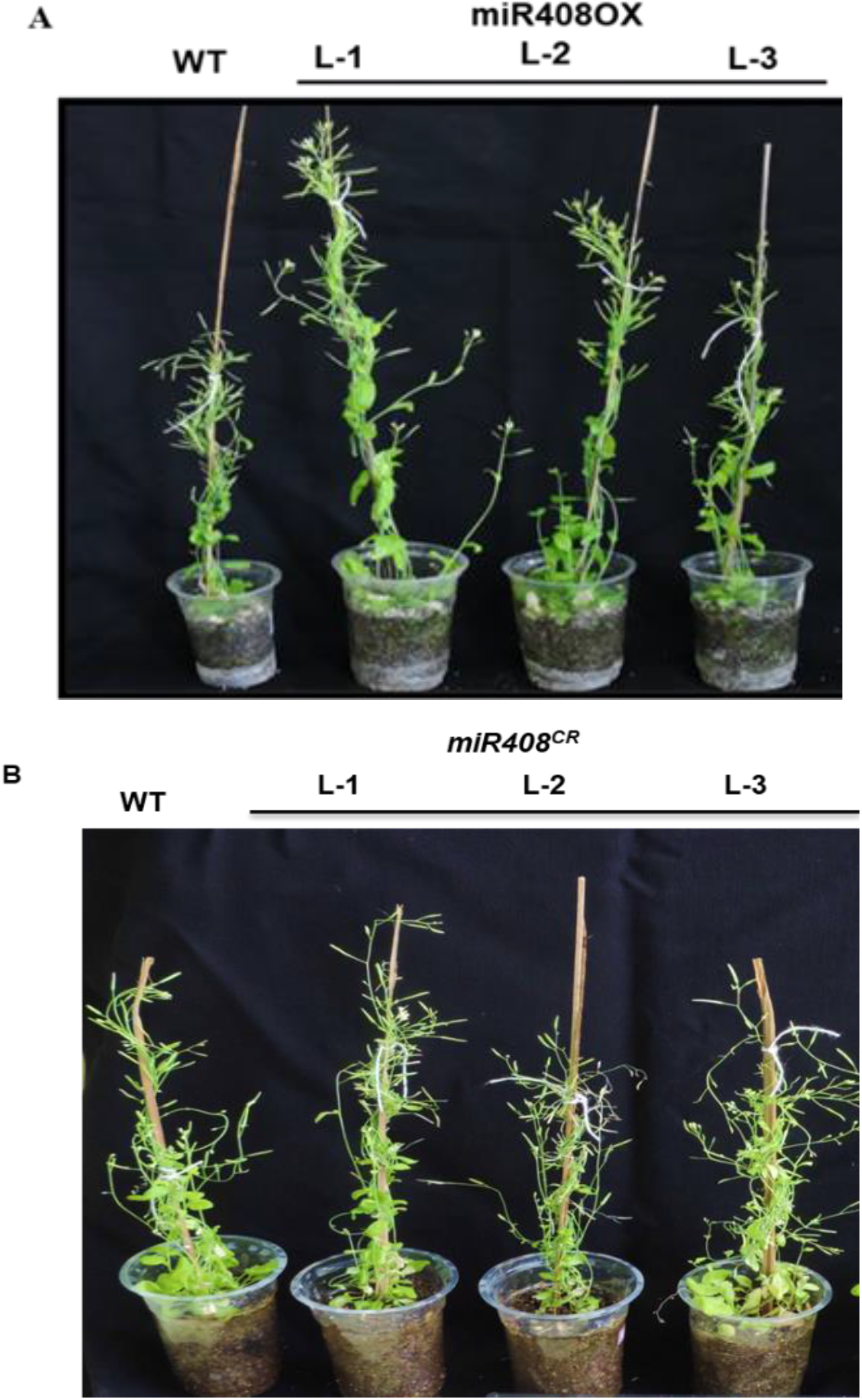
Representative image of mature WT, miR408 overexpressing and CRISPR-edited plants. **(A)** Image of miR408OX represented as L-1, L-2 and L-3 compare to the WT. **(B)** Image of miR408 CRISPR-edited plants represented as L-1, L-2 and L-3 compare to the WT (Scale bars,1 cm).

**Supplemental Figure S3.**
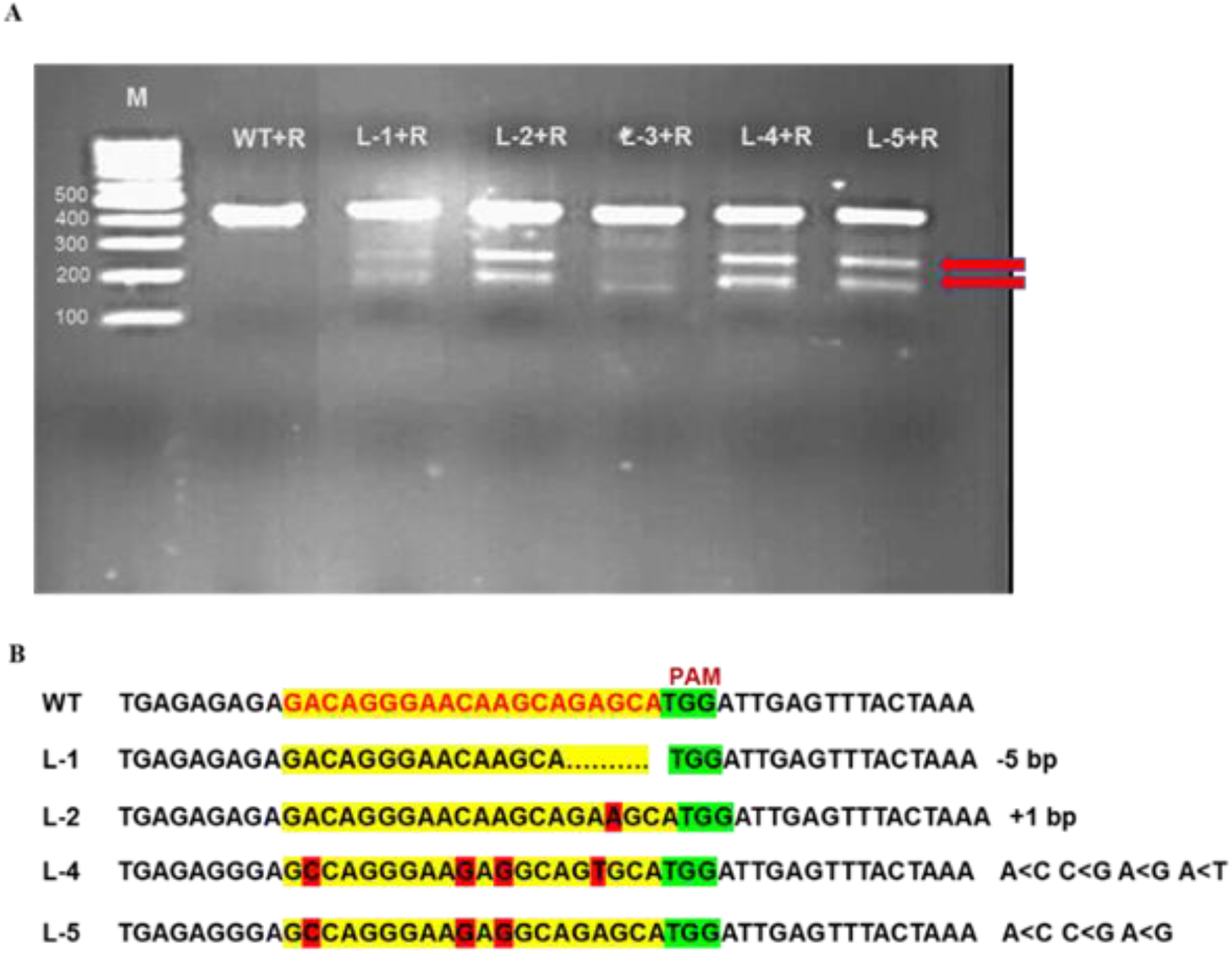
Analysis of CRISPR/Cas9 edited miR408 mutants. **(A)** For analysis of mutant lines, genomic DNA isolated from the *Arabidopsis* plants was transformed with pHSE401+gRNA and amplified by PCR with miPEP-specific primers. PCR product of WT and different lines were digested with resolvase enzyme (Takara). L-1 to L-5, are different edited plants with the addition of resolvase(+R). M, Molecular marker (100 bp ladder). (**B)** Analysis of sequence for target mutation in different plants (L-1, L-2, L-4 and L-5) compare to the WT. Deleted nucleotide showing as dots. The yellow highlight denotes the gRNA sequence used to target the sequence. Red highlights denote modified nucleotides through editing.

**Supplemental Figure S4.**
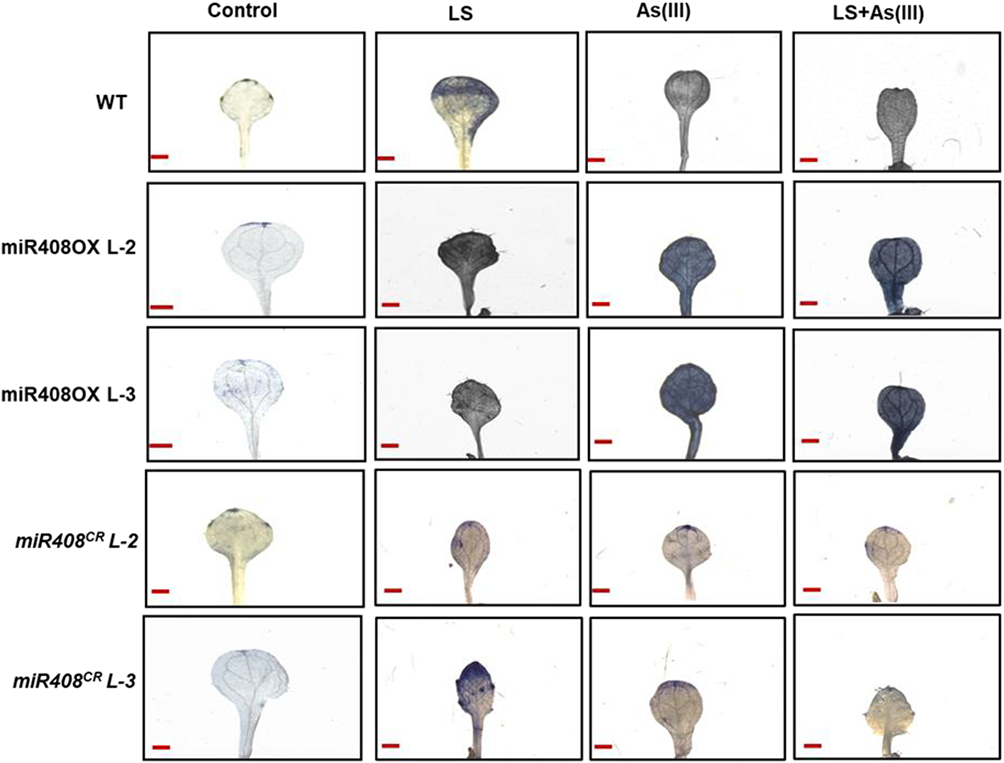
NBT staining in miR408OX and CRISPR-edited plants in limiting sulphur and As(III) stress. Staining of WT, miR408OX (L-2, and L-3) and *miR408^CR^* (L-2 and L-3) plants (L-2 and L-3) seedlings with Nitrotetrazolium (NBT) after grown for 10 days on medium containing optimum sulphur as control (C), limiting sulphur (LS), As (III) (10 μM), [LS + As(III)] (μM). Scale bar=1mm.

**Supplemental Figure S5.**
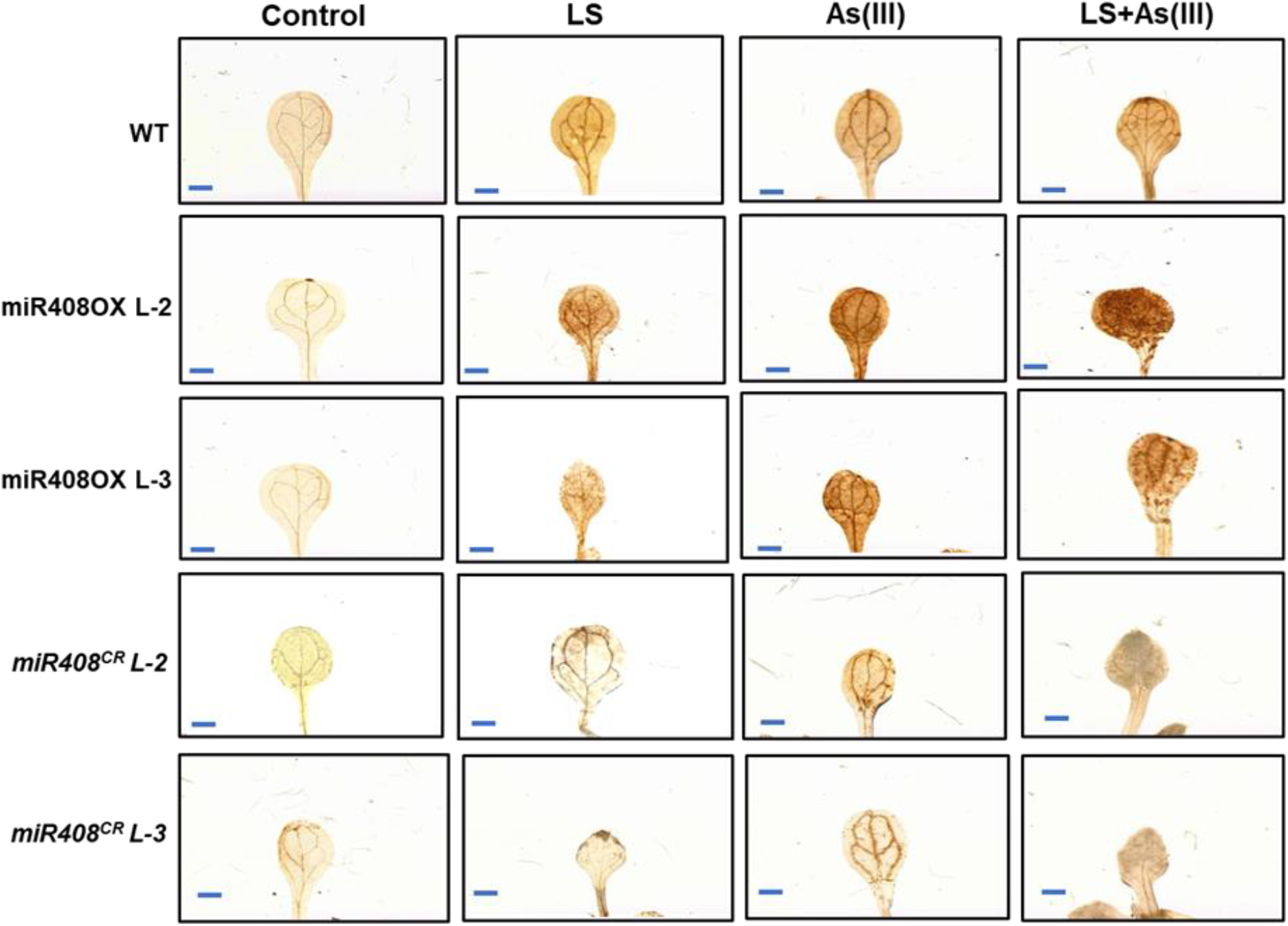
DAB staining in miR408OX and *miR408^CR^* plants in limiting sulphur and As(III) stress. Staining of WT, miR408OX (L-2, and L-3) and *miR408^CR^* plants (L-2 and L-3) seedlings with 3’-diaminobenzidine (DAB) after grown for 10 days on medium containing optimum sulphur as control (C), limiting sulphur (LS), As (III) (10 μM), [LS + As(III)] (10 μM). Scale bar= 1mm

**Supplemental Figure S6.**
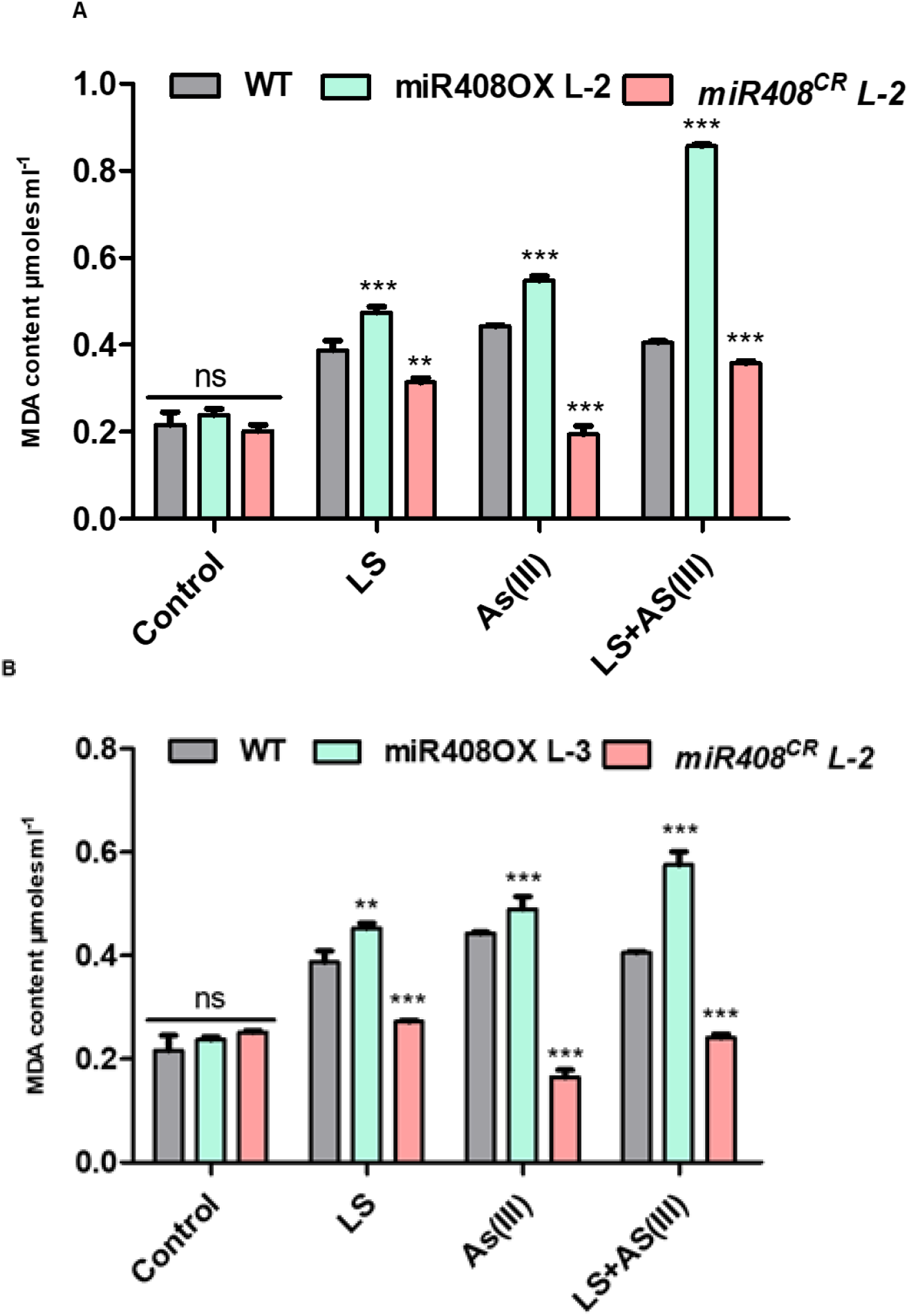
MDA content in miR408-OX and *miR408^CR^* seedlings. (A) Total MDA content of miR4080X (L-2) and CRISPR-edited *miR408^CR^* (L-2) ten-day-old seedlings under Control, LS, As(III) (10µM), [LS+As(III) (10 µM) (B) Total MDA content miR4080X (L-3) and CRISPR-edited *miR408^CR^* (L-3) ten-day old seedlings under Control, LS, As(III) (10µM), [LS+As(III) (10 µM). Calculations of data was performed from three biological replicated independently per treatment with similar results. The asterisk denotes significant difference in values with *** as P<0.001, according to two-tailed student’s t-test.

**Supplemental Figure S7.**
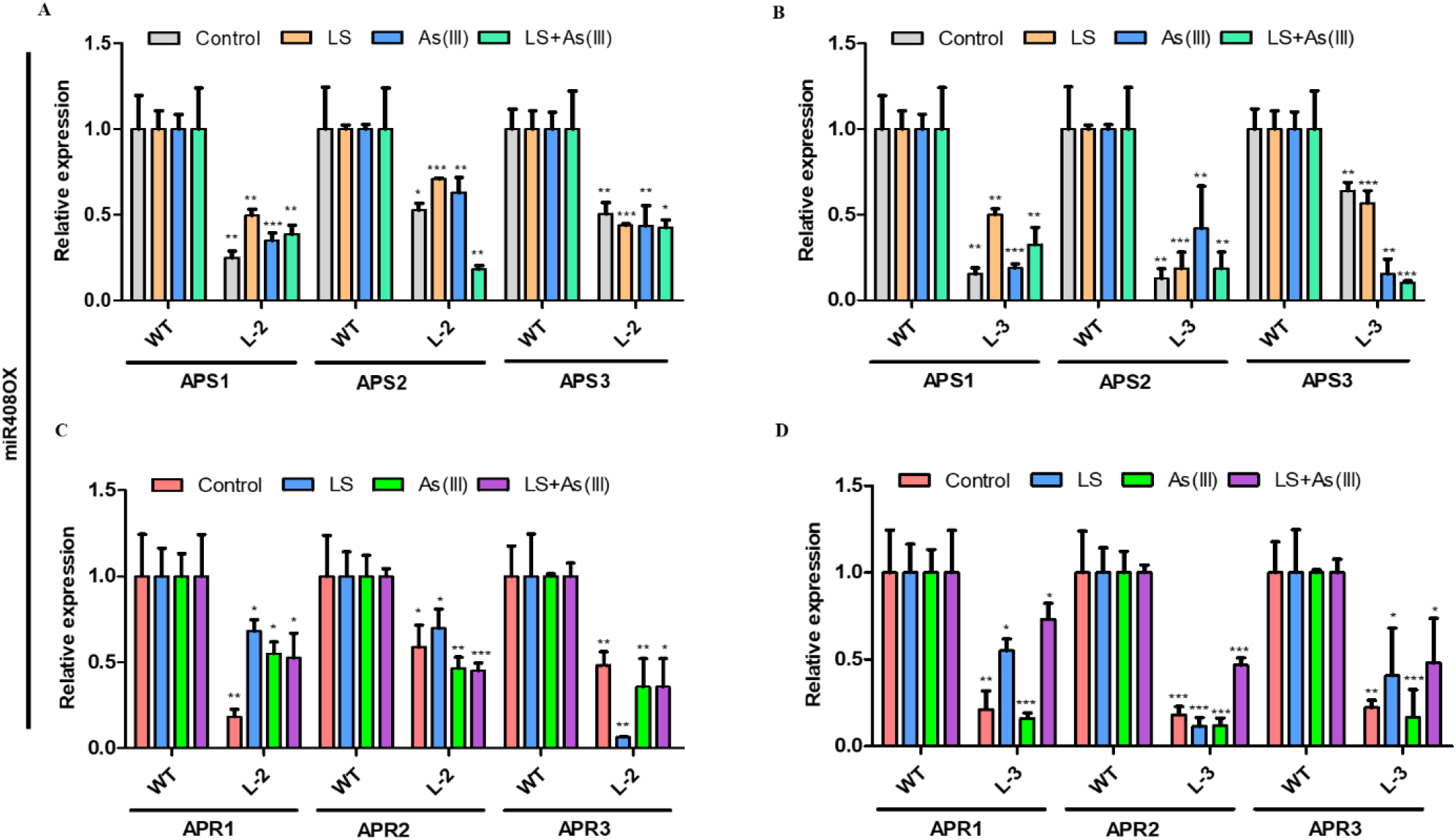
Relative expression of Sulphur reduction pathway genes in miR408-OX plants. **(A)** Relative expression of *AtAPS1*,*2* and *3* on miR408OX (L-2) plants under Control, LS, As(III) (10µM), [LS+As(III)] (10 µM) conditions. (**B)** Relative expression of *AtAPS1*,*2* and *3* on miR408-OX (L-3) plants under Control, LS, As(III) (10µM), [LS+As(III)] (10 µM). **C** Expression analysis of *AtAPR1*,2 and 3 in miR408OX (L-2) plants under Control, LS, As(III) (10µM), [LS+As(III)] (10 µM) conditions. (**D)** Expression analysis of *AtAPR1*,*2* and *3* in miR408-OX (L-3) plants under Control, LS, As(III) (10µM), [LS+As(III)] (10 µM) conditions. Calculations of data was performed from three biological replicated independently per treatment with similar results. The asterisk denotes significant difference in values with *** as *P < 0.1; **P < 0.01; ***P < 0.001, according to two-tailed student’s t-test.

**Supplemental Figure S8.**
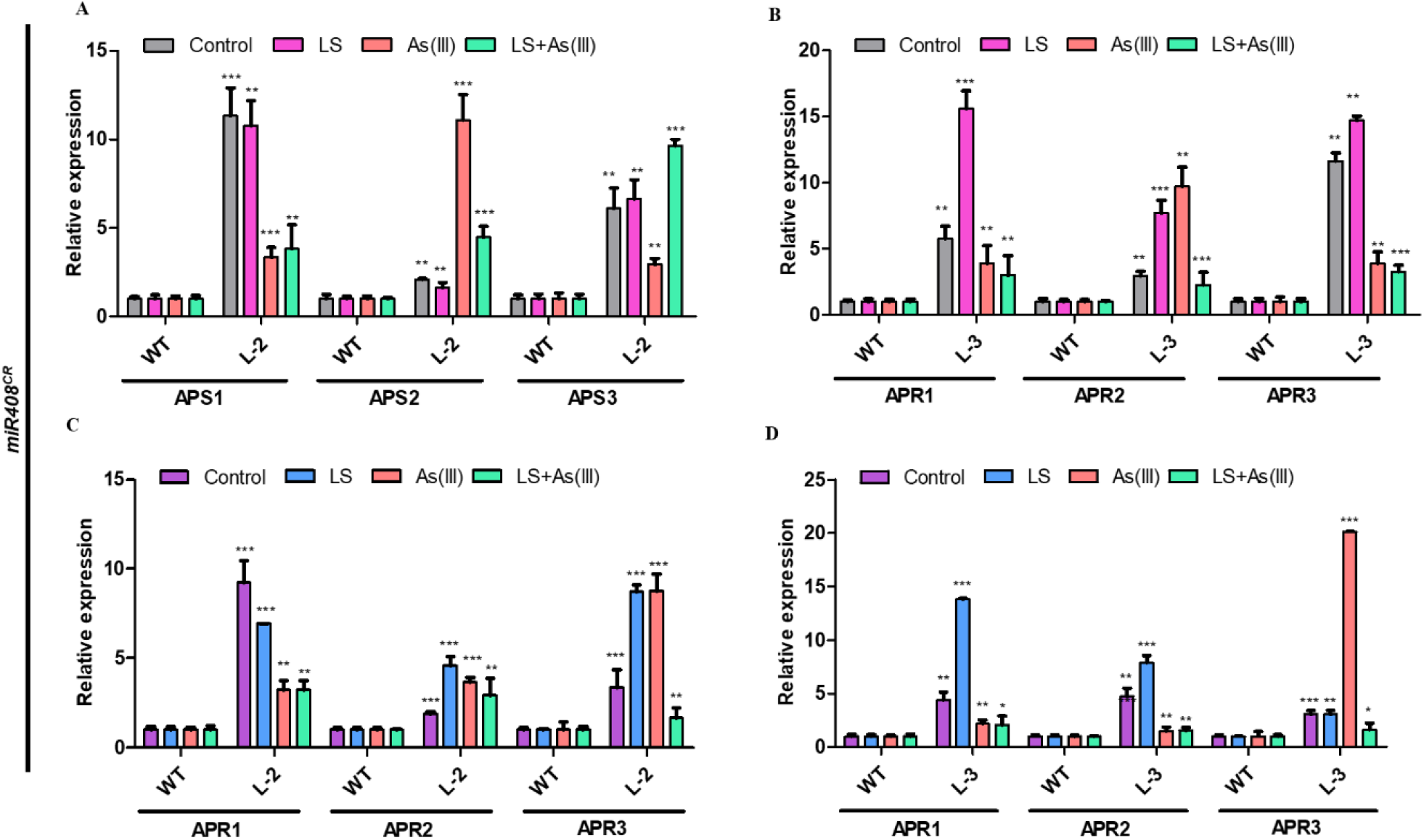
Relative expression of Sulphur reduction pathway genes in mutated *miR408^CR^* plants. **(A)** Relative expression of *AtAPS1*,2 and 3 on CRISPR-edited *miR408^CR^* (L-2) seedlings under Control, LS, As(III) (10µM), [LS+As(III)] (10 µM) conditions. (**B**) Relative expression of *AtAPS1*,*2* and *3* on CRISPR-edited *miR408^CR^* (L-3) plants under Control, LS, As(III) (10µM), [LS+As(III)] (10 µM) conditions. (**C)** Expression analysis of *AtAPR1*,2 and 3 in CRISPR-edited *miR408^CR^* (L-2) plants under Control, LS, As(III) (10µM), [LS+As(III)] (10 µM) conditions. (**D)** Expression analysis of *AtAPR1*,*2* and *3* in CRISPR-edited *miR408^CR^* (L-3) plants under Control, LS, As(III) (10µM), [LS+As(III)] (10 µM) conditions. Data are mean ±SD calculated from three biological replicated. *** represent show significant from control at P < 0.001, according to student’s t-test.

**Supplemental Figure S9.**
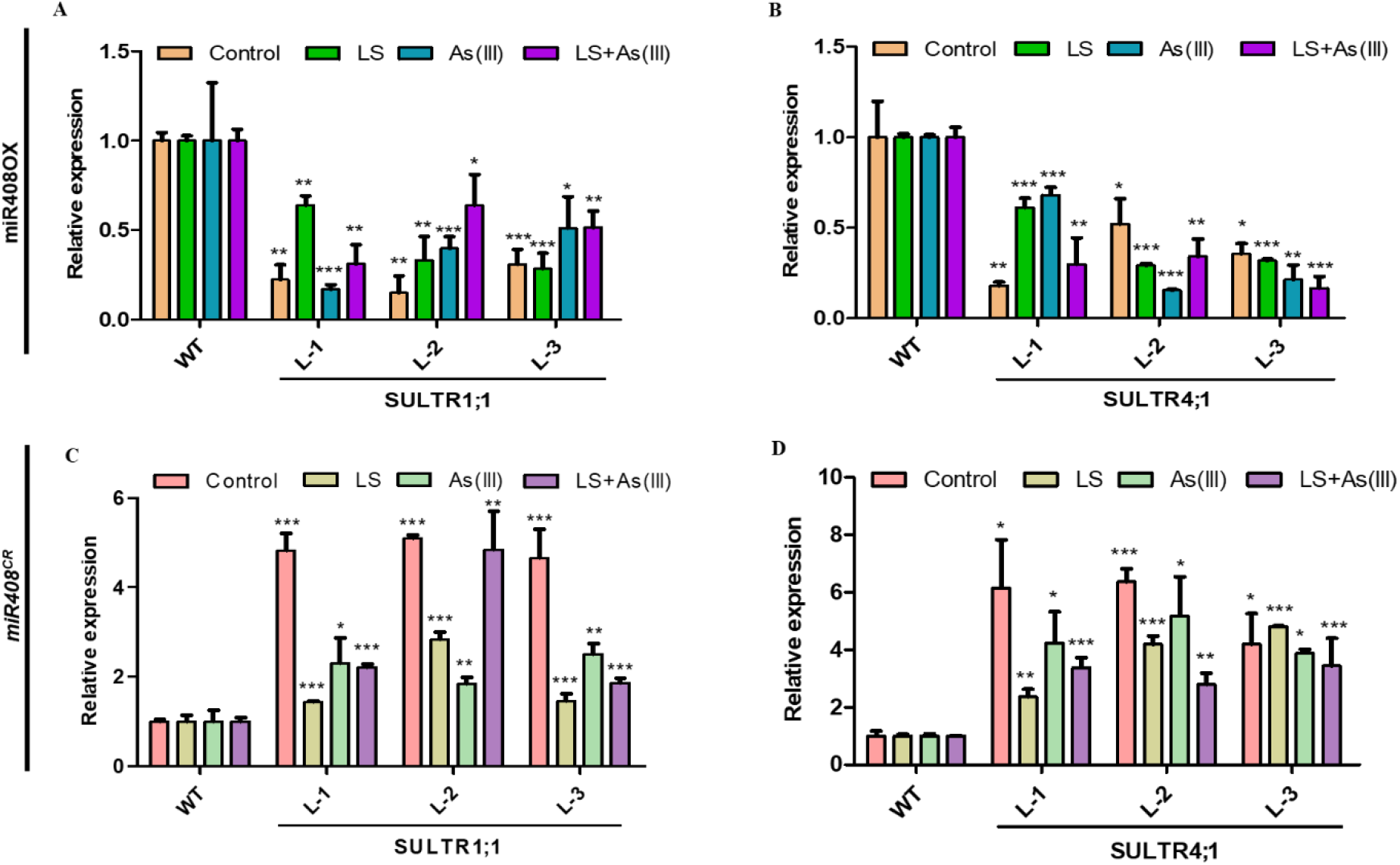
Relative expression of SULTR1;1 and SULTR4;1 in miR408-OX and edited *miR408^CR^* plants. **(A)** Relative expression of *SULTR1;1* in miR408OX (L1, L2 and L-3) ten-day old seedlings under Control, LS, As(III) (10µM), [LS+As(III)] (10 µM) conditions. (**B**) Relative expression of *SULTR4;1* in miR408-OX (L-1, L-2 and L-3) ten-day old seedlings under Control, LS, As(III) (10µM), [LS+As(III)] (10 µM) conditions. (**C)** Expression analysis of *SULTR 1;1* in CRISPR-edited *miR408^CR^* (L-1, L-2 and L-3) plants under Control, LS, As(III) (10µM), [LS+As(III)] (10 µM) conditions. (**D)** Expression analysis of *SULTR 4;1* in CRISPR-edited *miR408^CR^* (L-1, L-2 and L-3) plants under Control, LS, As(III) (10µM), [LS+As(III)] (10 µM) conditions. Data are mean ±SD calculated from three biological replicated. *** represent show significant from control at P < 0.001, according to student’s t-test.

**Supplemental Figure S10.**
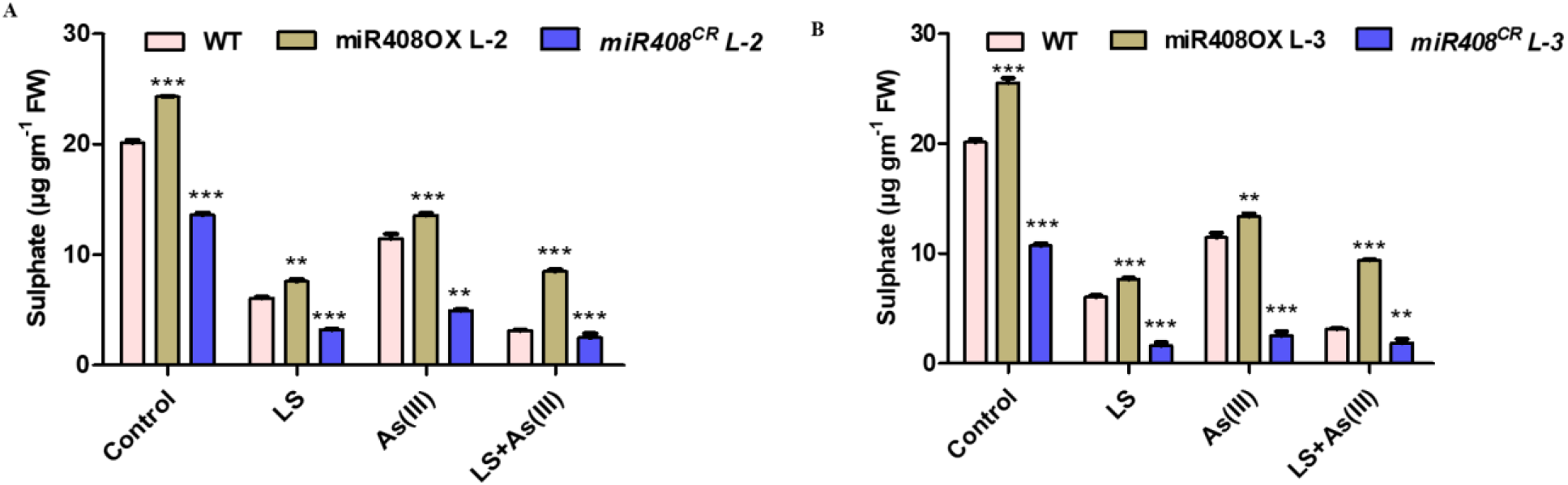
Total sulphur content in miR408-OX and edited *miR408^CR^* plants. **(A)** Total sulphur content of miR4080X (L-2) and CRISPR-edited *miR408^CR^* (L-2) 10 days old seedlings under Control, LS, As(III) (10µM), [LS+As(III) (10 µM) conditions. **(B)** Total sulphur content of miR4080X (L-3) and CRISPR-edited *miR408^CR^* (L-3) ten-day old seedlings under Control, LS, As(III) (10µM), [LS+As(III) (10 µM) conditions. Calculations of data was performed from three biological replicated independently per treatment with similar results. The asterisk denotes significant difference in values with *** as *P < 0.1; **P < 0.01; ***P < 0.001, according to two-tailed student’s t-test.

**Supplemental Figure S11.**
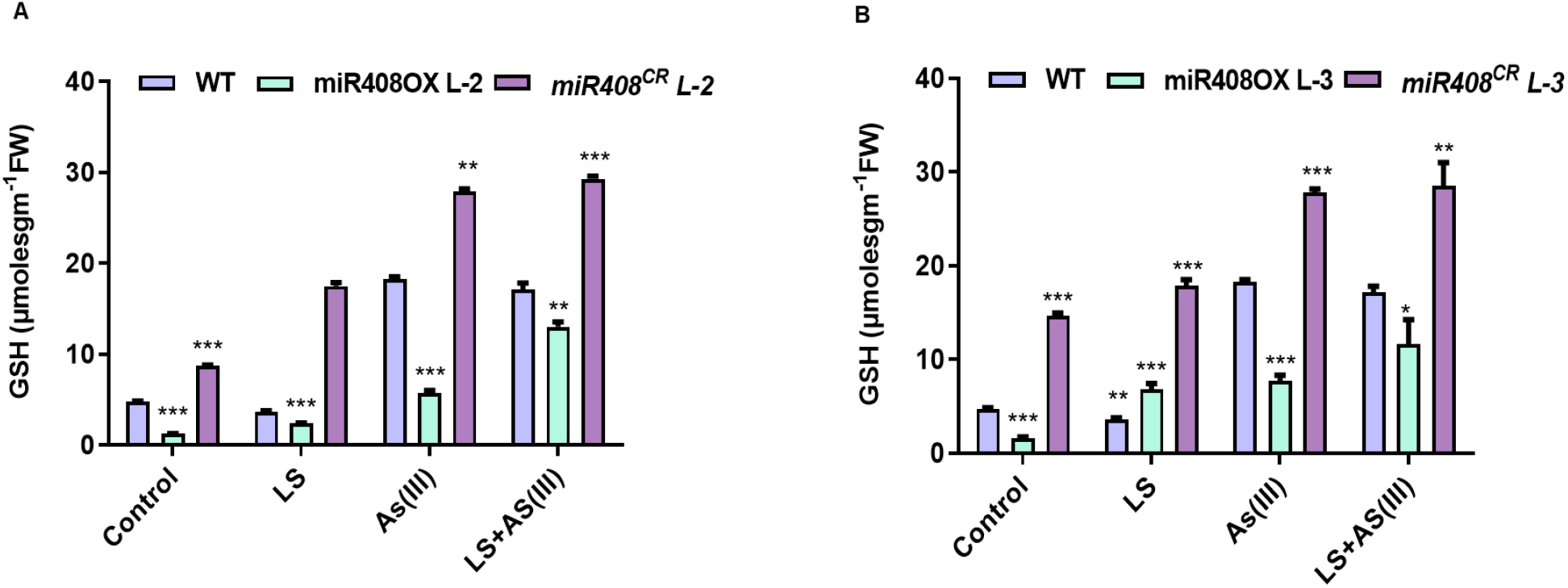
Total glutathione (GSH) estimation in miR408-OX and *mir408^CR^* plants. **(A)** Total GSH content of miR4080X (L-2) and CRISPR-edited *miR408^CR^* (L-2) 10 days old seedlings under Control, LS, As(III) (10µM), [LS+As(III) (10 µM) conditions. **(B)** Total GSH content of miR4080X (L-3) and CRISPR-edited *miR408^CR^* (L-3) ten-day old seedlings under Control, LS, As(III) (10µM), [LS+As(III) (10 µM) conditions. Calculations of data was performed from three biological replicated independently per treatment with similar results. The asterisk denotes significant difference in values with *** as *P < 0.1; **P < 0.01; ***P < 0.001, according to two-tailed student’s t-test.

**Supplemental Figure S12.**
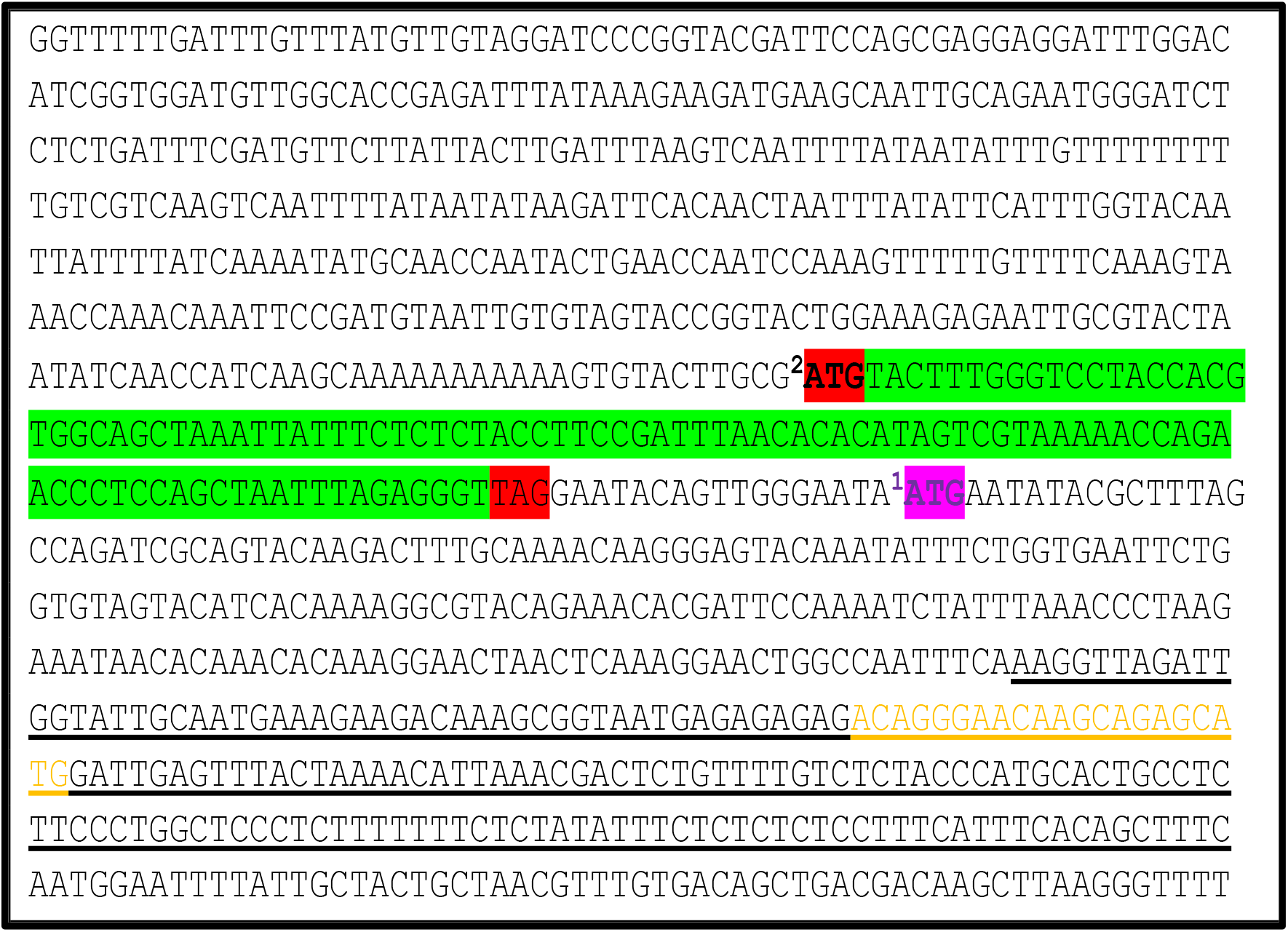
Upstream sequence (1000 bp) from pre-mi408 represents putative ORFs. ATG1 and ATG2 are the start codon of putative ORFs. The ORF encoding functional miPEP408 peptide is the green highlight. The underlined sequence represents the pre-miR408 sequence. The mature miR408 sequence is shown in yellow colour.

**Supplemental Figure S13.**
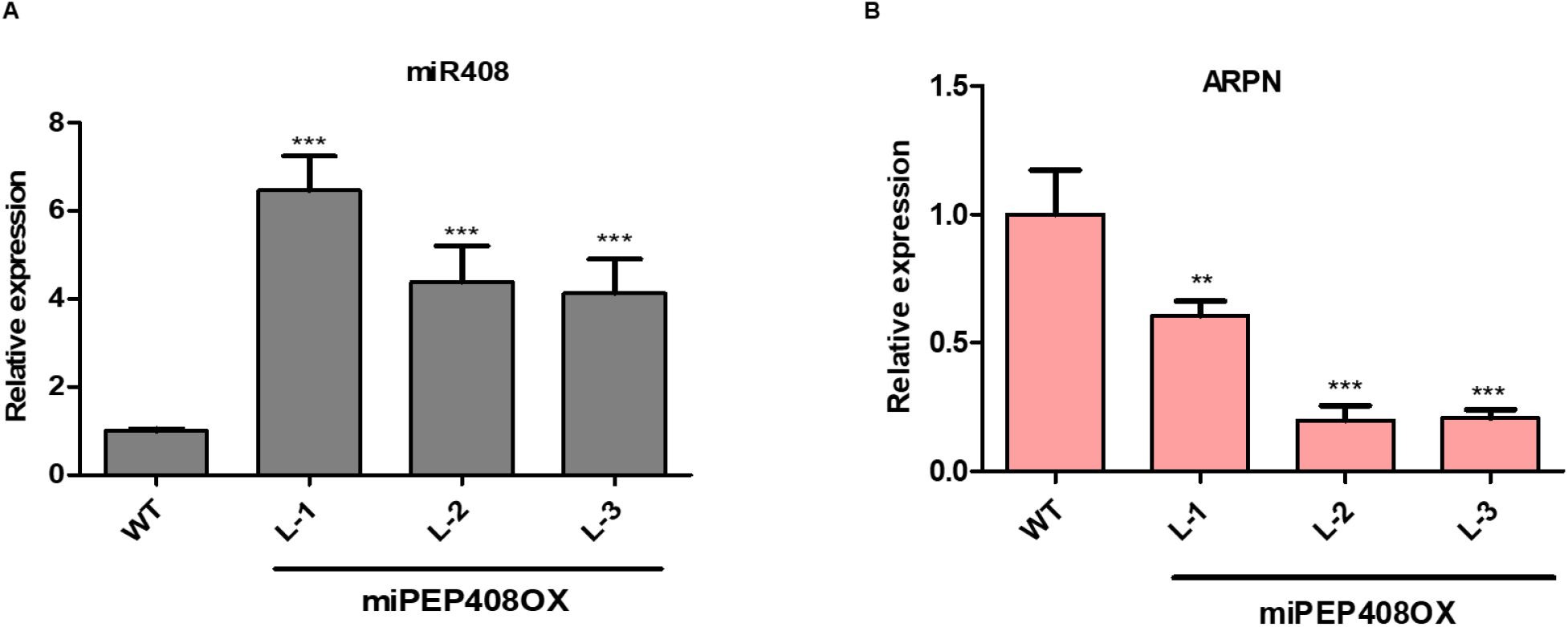
Expression analysis of miR408 and its target in miPEPOX plants. **(A)** Expression analysis of pre-miR408 in 10 days old WT and miPEP408-OX plants. Three independent experiments were performed, with similar results. (**B**) Relative expression of *ARPN* transcript in 10 days old WT and miR408 overexpressing plants. Calculations of data were performed from three biological replicated independently per treatment with similar results. The asterisk denotes significant difference in values with *** as *P < 0.1; **P < 0.01; ***P < 0.001, according to two-tailed student’s t-test.

**Supplemental Figure S14.**
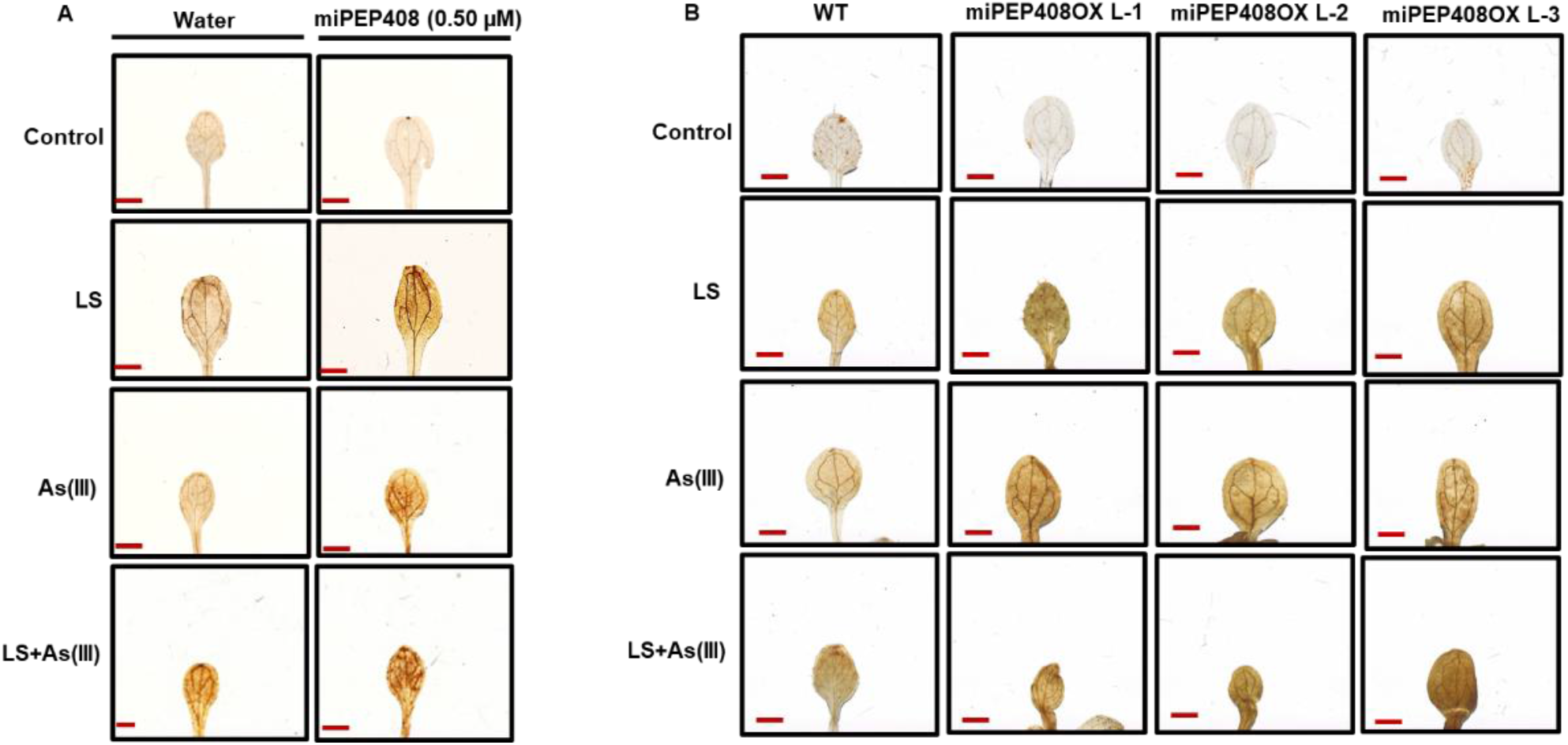
DAB staining of Exogenous miPEP408 supplemented and miPEP408OX 10 days old seedlings grown on limiting sulphur and As(III) stress. **(A)** 3-3’diaminobenzidine (DAB) staining of WT *Arabidopsis* seedlings grown for 10 days on control and stress conditions supplemented with Water and miPEP408 separately. **(B)** Staining of WT and miPEP408-OX seedlings with 3’-diaminobenzidine (DAB) after grown for 10 days on medium containing optimum sulphur as control (C), limiting sulphur (LS), As (III) (10 μM), [LS + As(III)] (10 μM). Scale bar= 1mm.

**Supplemental Figure S15.**
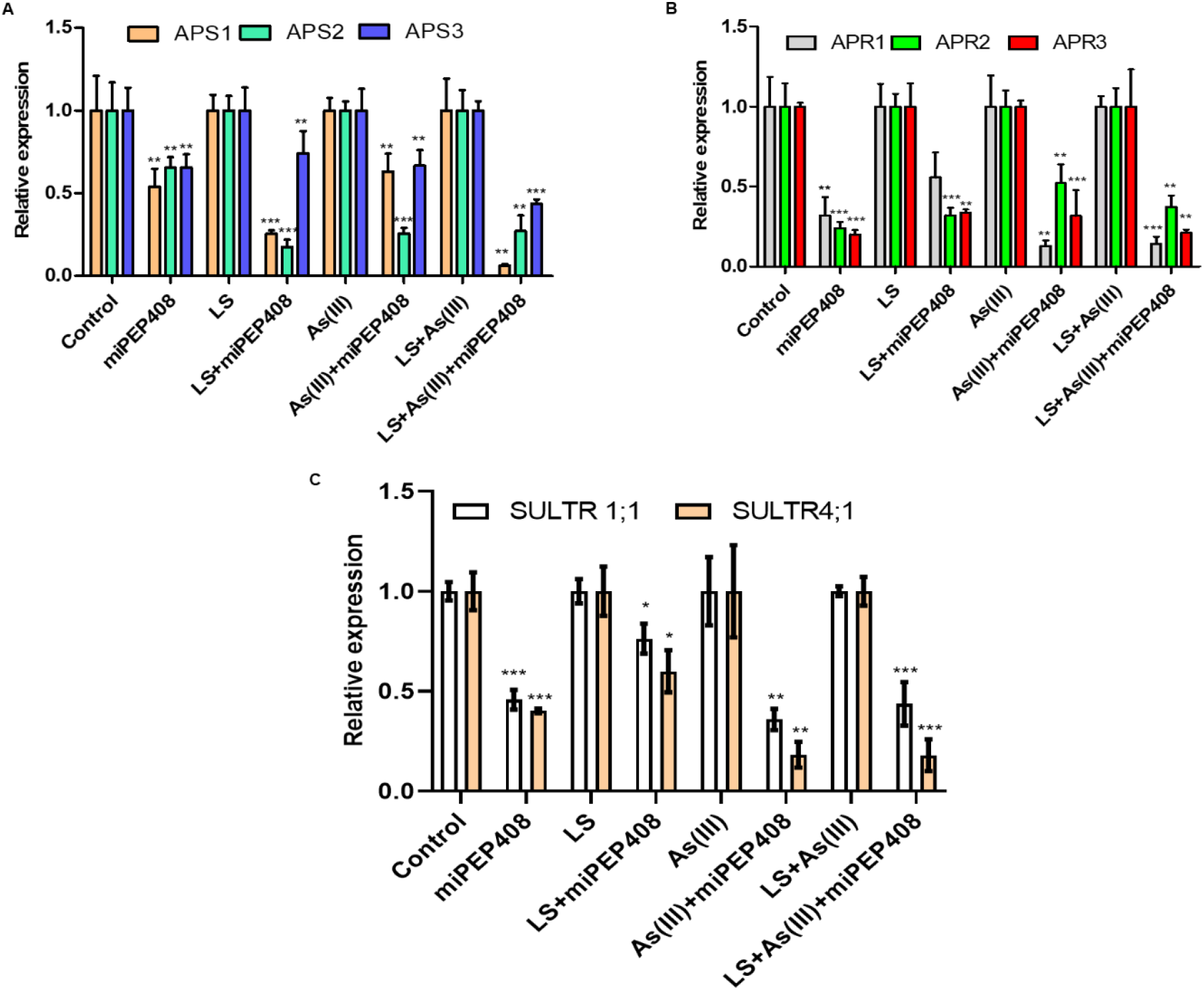
miPEP408 regulate sulphur metabolism and affect plant tolerance toward stress. **(A)** Relative expression of *AtAPS1*,2 and *3* in ten days grown WT *Arabidopsis* seedlings grown under Control, LS, As(III) (10µM), [LS+As(III)] (10 µM) conditions supplemented with water and miPEP408 individually. (**B)** Relative expression of *AtAPR1*,2 and *3* on ten days grown WT *Arabidopsis* seedlings under Control, LS, As(III) (10µM), [LS+As(III)] (10 µM) conditions supplemented water and miPEP408 individually. (**C)** The expression on SULTR1;1 and SULTR4;1of ten-day grown seedlings on media containing Control, LS, As(III) and [LS+As(III)]. Calculations of data were performed from three biological replicated independently per treatment with similar results. The asterisk denotes significant difference in values with *** as *P < 0.1; **P < 0.01; ***P < 0.001, according to two-tailed student’s t-test.

**Supplemental Figure S16.**
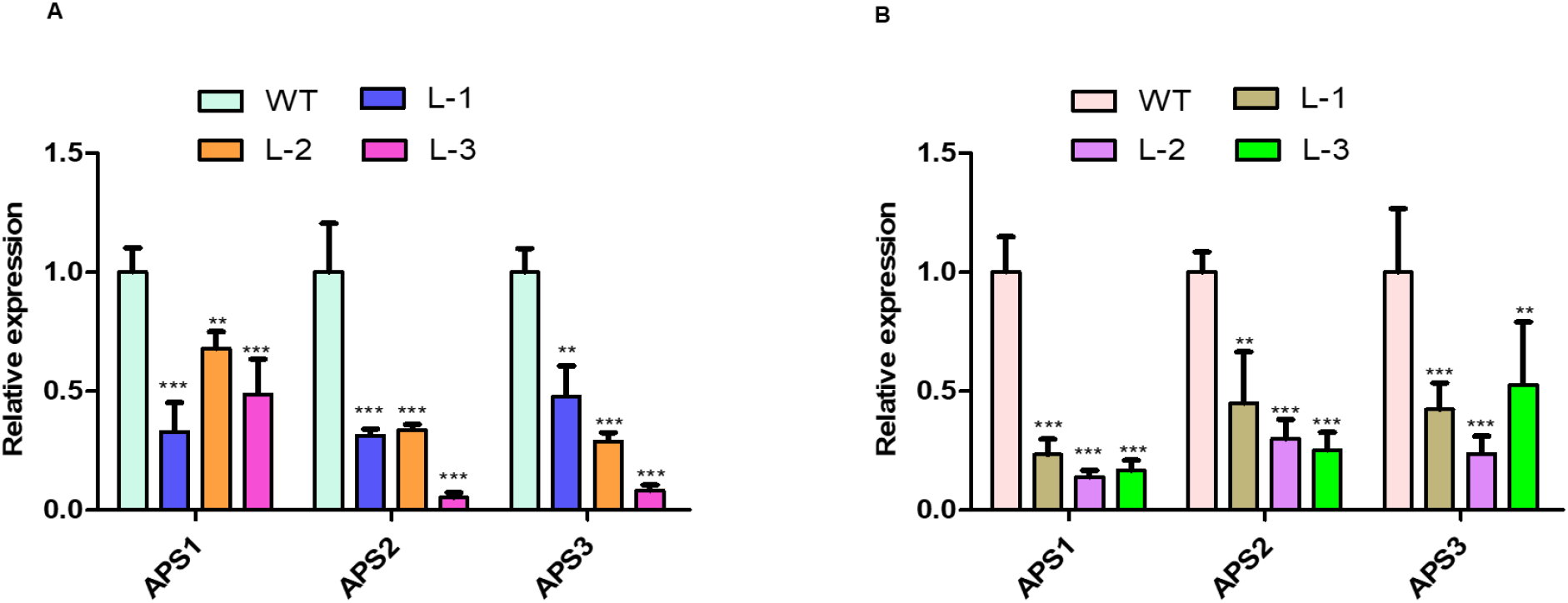
Overexpression of miPEP408 regulates sulphur assimilation pathway and plant stress response. Relative expression of *AtAPS1*,2 and *3* in ten days grown WT *Arabidopsis* seedlings grown under control conditions (**B)** Relative expression of *AtAPR1*,2 and *3* in ten days grown WT *Arabidopsis* seedlings under Control conditions supplemented water and miPEP408 individually. Calculations of data were performed from three biological replicated independently per treatment with similar results. The asterisk denotes significant difference in values with *** as *P < 0.1; **P < 0.01; ***P < 0.001, according to two-tailed student’s t-test.

**Supplemental Figure S17.**
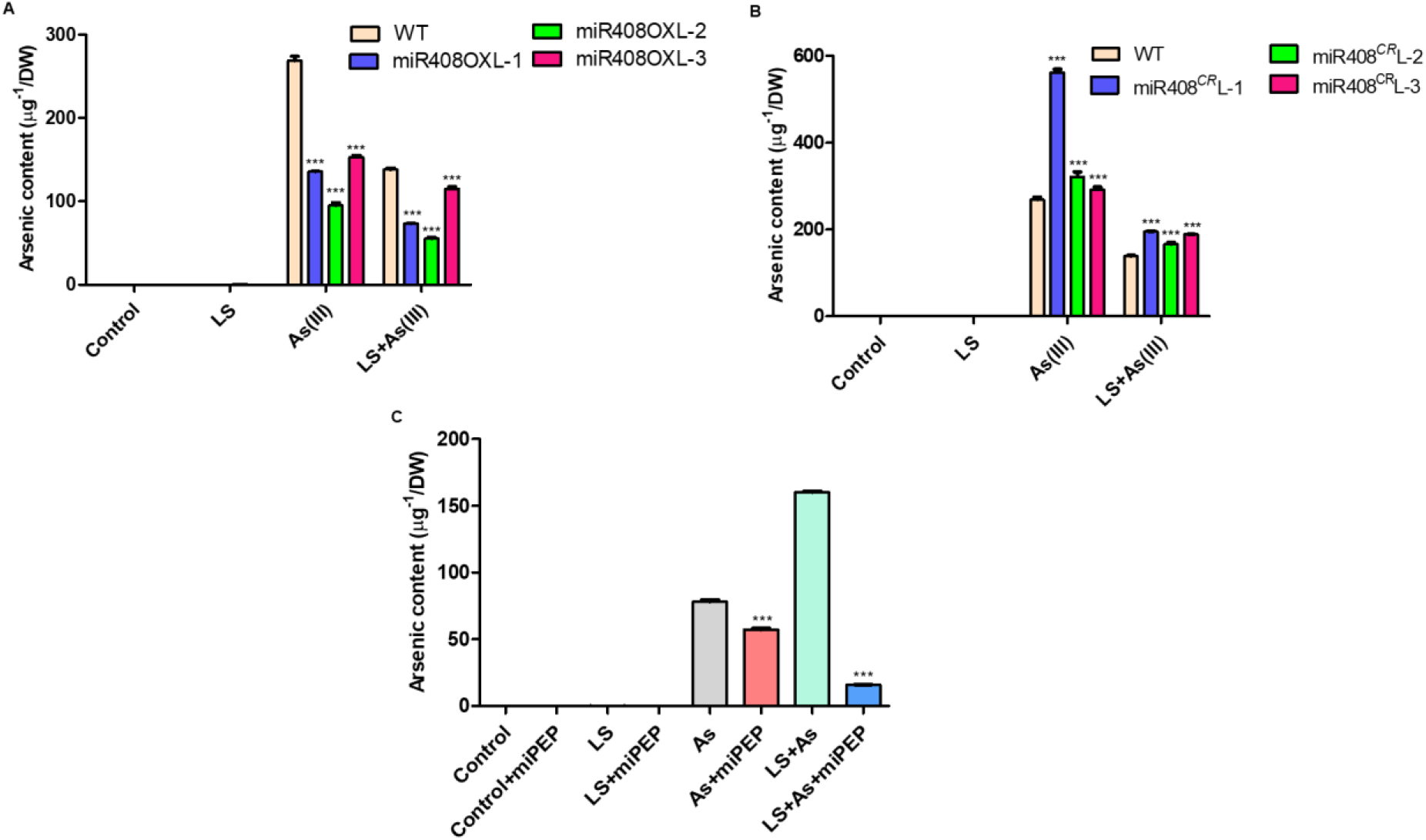
miR408 affects the arsenic accumulation leading to the sensitivity and tolerance phenotype of Arabidopsis seedlings in heavy metal stress. (A) Arsenic estimation using ICP-MS in the miR408-OX compared to the WT (B) Arsenic estimation using ICP-MS in the *miR408^CR^* compared to the WT (C) Arsenic estimation of COL-O 10d days old seedlings supplemented with peptide in different stress conditions. Three biological replicates per treatment are taken to calculate mean±SD. ∗∗∗ represent values that differ significantly from control at P<0.001, according to the student’s unpaired t-test.

**Supplemental Table S1.** The small RNA sequencing summary.

| S. No. | Sample | Total Reads<br>(Single-End) | Sequence Length<br>(bp) | Data<br>(Mb) | %GC | Phred<br>score | SRA details |
| --- | --- | --- | --- | --- | --- | --- | --- |
| 1. | AT-CON | 44,749,679 | 50 | 2237.48 | 51.07 | 39.95 | SRX14947982 |
| 2. | AT-LS | 42,343,343 | 50 | 2117.17 | 52.61 | 40.0 | SRX14947983 |
| 3. | AT-ASIII | 44,650,101 | 50 | 2232.51 | 52.72 | 39.97 | SRX14947984 |
| 4. | AT-LS+ASIII | 71,641,633 | 50 | 3582.08 | 52.99 | 39.99 | SRX14947985 |

**Supplemental Table S2.** Overall summary of raw reads of samples after adapter removal.

| **S. No.** | **Sample name** | **Total number of raw reads** | **Number of unique reads (after adapter removal)** |
| --- | --- | --- | --- |
| **1.** | AT-CON | 44,749,679 | 6,933,010 |
| **2.** | AT-LS | 42,343,343 | 5,459,037 |
| **3.** | AT-ASIII | 44,650,101 | 5,010,037 |
| **4.** | AT-LS+ASIII | 71,641,633 | 8,340,079 |

**Supplemental Table S3.** Summary of known miRNA predicted from samples with the reference genome.

| S. No. | Samples | No. of known miRNAs |
| --- | --- | --- |
| 1. | AT-CON | 14280 |
| 2. | AT-LS | 10317 |
| 3. | AT-ASIII | 7868 |
| 4. | AT-LS+ASIII | 15468 |

**Supplemental Table S4.** Differentially expressed miRNAs under different stress conditions.

| miRNAs | Ct_vs_Ls | Ct_vs_As | Ct_vs_Ls+As |
| --- | --- | --- | --- |
| ath-miR398a-3p | -7.398521 | -5.7310881 | -5.6911619 |
| ath-MIR398b | -6.798502 | -5.686116 | -7.0355082 |
| ath-miR398b-3p | -7.6266785 | -6.2654064 | -7.0079209 |
| ath-miR398b-5p | -5.3058221 | -4.7676721 | -4.5333679 |
| ath-MIR398c | -6.7976915 | -5.6853054 | -7.0653944 |
| ath-miR398c-3p | -7.6266785 | -6.2654064 | -7.0079209 |
| ath-miR398c-5p | -5.3058221 | -4.7676721 | -4.5333679 |
| ath-MIR399a | -9.4177241 | -7.1426085 | -7.6738391 |
| ath-miR399b | -4.1969711 | -3.6099116 | -3.4431706 |
| ath-MIR399c | -5.2293118 | -4.9541963 | -5.8073549 |
| ath-miR399c-3p | -4.1969711 | -3.6099116 | -3.4431706 |
| ath-miR399c-5p | -6.8649004 | -5.5897849 | -7.4429435 |
| ath-MIR399d | -7.8976903 | -7.6225748 | -8.4757334 |
| ath-MIR399f | -8.485352 | -7.2102365 | -7.4784326 |
| ath-MIR408 | -7.1447644 | -5.2846864 | -6.4008794 |
| ath-miR408-3p | -4.2014607 | -4.6730588 | -3.6539729 |
| ath-miR408-5p | -4.9117401 | -5.1313893 | -4.3146965 |
| ath-miR857 | -4.8813885 | -4.606273 | -3.4594316 |
| ath-MIR169d | -2.3737017 | 3.09642976 | 0 |
| ath-MIR169e | -4.8142743 | 3.26174109 | 0 |
| ath-miR169f-3p | 0 | 2.97042466 | 0 |
| ath-MIR169g | -2.3737017 | 3.09642976 | 0 |
| ath-miR2111b-3p | 0 | 0 | -4.6438562 |
| ath-MIR391 | 0 | 0 | -3.0914984 |
| ath-miR391-3p | 0 | 0 | -2.5168735 |
| ath-miR391-5p | 0 | 0 | -2.7369656 |
| ath-MIR395a | 6.9866427 | 0 | 7.8964104 |
| ath-MIR395b | 7.597745 | 0 | 6.0435195 |
| ath-MIR395c | 7.5937369 | 0 | 6.0426443 |
| ath-MIR395d | 9.4282299 | 0 | 8.8845868 |
| ath-MIR395e | 6.9659499 | 0 | 7.8964104 |
| ath-MIR395f | 9.5541776 | 0 | 7.0465784 |
| ath-miR397b | 0 | -4.6704033 | -5.523562 |
| ath-MIR398a | -5.3992368 | 0 | 0 |
| ath-miR398a-5p | 0 | 0 | 3.2854022 |
| ath-miR399e | -5.3526943 | 0 | -5.9307373 |
| ath-miR5996 | 0 | 0 | -3.3923174 |
| ath-miR8167a | -5.0943823 | -4.8192667 | 0 |
| ath-miR8167b | -5.0943823 | -4.8192667 | 0 |
| ath-miR8167c | -5.0943823 | -4.8192667 | 0 |
| ath-miR8167d | -5.0943823 | -4.8192667 | 0 |
| ath-miR8167e | -5.0943823 | -4.8192667 | 0 |
| ath-miR8167f | -5.0943823 | -4.8192667 | 0 |
| ath-MIR8171 | -2.662965 | 0 | 0 |
| ath-MIR8173 | 0 | -3.2726853 | 0 |
| ath-miR8176 | 0 | 0 | 4.9068906 |
| ath-miR826a | -2.8387967 | -4.4001824 | 0 |
| ath-miR827 | -4.0702983 | 0 | -3.5891022 |
| ath-miR828 | 4.4525122 | 0 | 6.3692338 |
| ath-miR845b | 0 | 0 | 4.0297473 |
| ath-miR858a | 0 | 0 | 2.7104934 |
| ath-MIR858b | 0 | 0 | 3.1858665 |
| ath-miR866-5p | 2.7185243 | 0 | 0 |
| ath-MIR869 | 0 | 0 | 2.5326473 |
| ath-miR869.2 | 0 | 0 | 2.7151612 |

**Supplemental Table S5:**
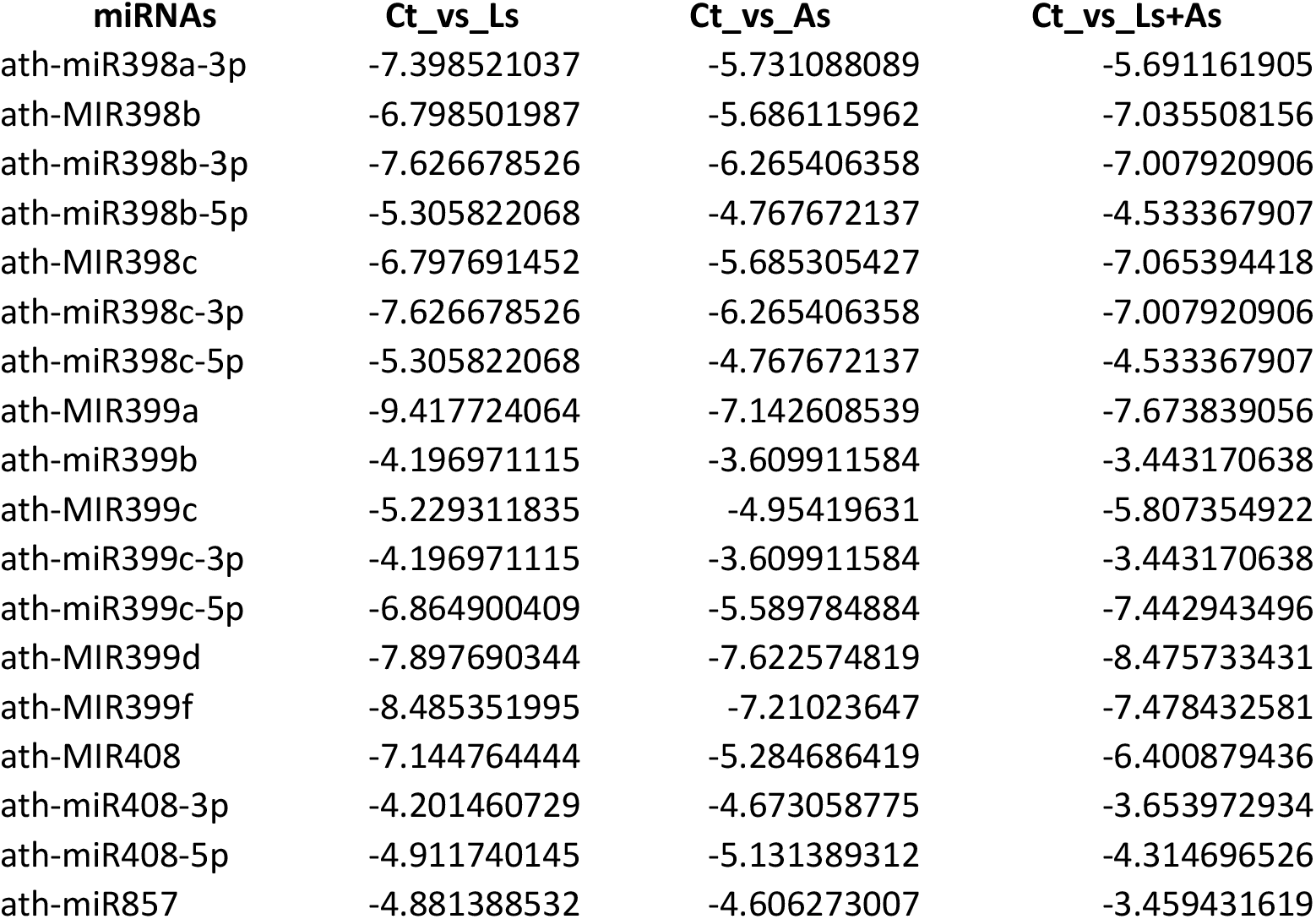
Commonly differentially expressed miRNAs under different stress conditions.

| **miRNAs** | **Ct_vs_Ls** | **Ct_vs_As** | **Ct_vs_Ls+As** |
| --- | --- | --- | --- |
| ath-miR398a-3p | -7.398521037 | -5.731088089 | -5.691161905 |
| ath-MIR398b | -6.798501987 | -5.686115962 | -7.035508156 |
| ath-miR398b-3p | -7.626678526 | -6.265406358 | -7.007920906 |
| ath-miR398b-5p | -5.305822068 | -4.767672137 | -4.533367907 |
| ath-MIR398c | -6.797691452 | -5.685305427 | -7.065394418 |
| ath-miR398c-3p | -7.626678526 | -6.265406358 | -7.007920906 |
| ath-miR398c-5p | -5.305822068 | -4.767672137 | -4.533367907 |
| ath-MIR399a | -9.417724064 | -7.142608539 | -7.673839056 |
| ath-miR399b | -4.196971115 | -3.609911584 | -3.443170638 |
| ath-MIR399c | -5.229311835 | -4.95419631 | -5.807354922 |
| ath-miR399c-3p | -4.196971115 | -3.609911584 | -3.443170638 |
| ath-miR399c-5p | -6.864900409 | -5.589784884 | -7.442943496 |
| ath-MIR399d | -7.897690344 | -7.622574819 | -8.475733431 |
| ath-MIR399f | -8.485351995 | -7.21023647 | -7.478432581 |
| ath-MIR408 | -7.144764444 | -5.284686419 | -6.400879436 |
| ath-miR408-3p | -4.201460729 | -4.673058775 | -3.653972934 |
| ath-miR408-5p | -4.911740145 | -5.131389312 | -4.314696526 |
| ath-miR857 | -4.881388532 | -4.606273007 | -3.459431619 |

**Supplemental Table S6.**
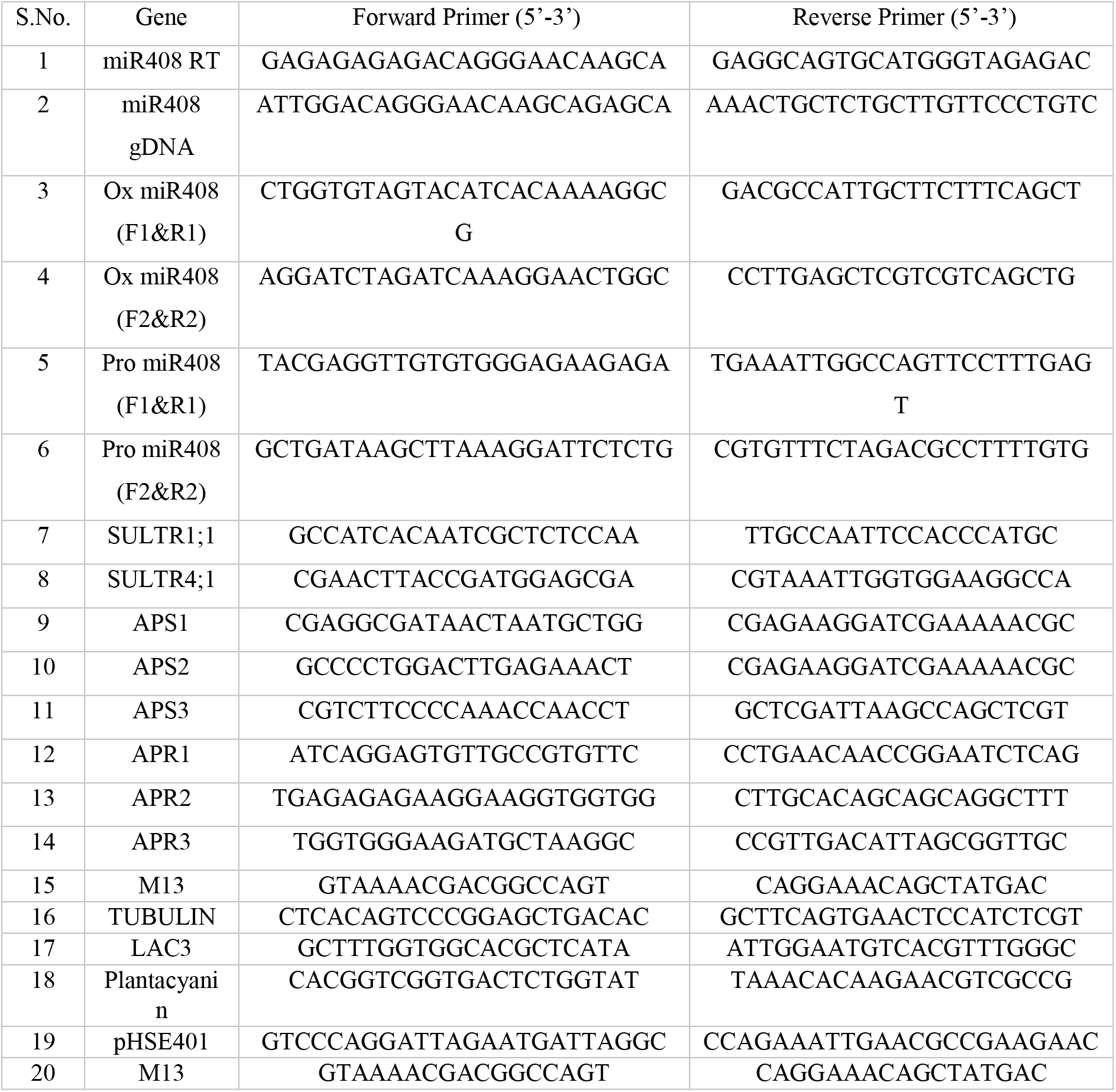
Oligonucleotides used for the development of constructs and expression analysis.

